# Separation of sticker-spacer energetics governs the coalescence of metastable condensates

**DOI:** 10.1101/2023.10.03.560747

**Authors:** Aniruddha Chattaraj, Eugene I. Shakhnovich

## Abstract

Biological condensates often emerge as a multi-droplet state and never coalesce into one large droplet within the experimental timespan. Previous work revealed that the sticker-spacer architecture of biopolymers may dynamically stabilize the multi-droplet state. Here, we simulate the condensate coalescence using metadynamics approach and reveal two distinct physical mechanisms underlying the fusion of droplets. Condensates made of sticker-spacer polymers readily undergo a kinetic arrest when stickers exhibit slow exchange while fast exchanging stickers at similar levels of saturation allow merger to equilibrium states. On the other hand, condensates composed of homopolymers fuse readily until they reach a threshold density. Increase in entropy upon inter-condensate mixing of chains drives the fusion of sticker-spacer chains. We map the range of mechanisms of kinetic arrest from slow sticker exchange dynamics to density mediated in terms of energetic separation of stickers and spacers. Our predictions appear to be in qualitative agreement with recent experiments probing dynamic nature of protein-RNA condensates.

**Statement of significance:** A key conundrum of biological condensates is the coexistence of multiple droplets, in direct variance with classical predictions of mean-field theories of polymer solutions. Our current study uncovers that the merging of sticker-spacer condensate is an entropy driven process, as opposed to the surface energy driven fusion that are observed for canonical liquid droplets. This entropy, stemming from the inter-condensate polymer exchange, makes the droplet merging process dependent on inter-sticker dissociation kinetics. Stronger inter-sticker interaction triggers a kinetic arrest, preventing the condensate merger even at a low density. Our prediction starkly correlates with recent experimental findings on protein-RNA condensates in vitro and in vivo, highlighting the biological relevance of the interplay of kinetics and thermodynamics.

## Introduction

Biomolecular condensates are membrane-less intracellular compartments that emerge via phase transition (1, 2). Condensates perform a range of spatiotemporal biochemical tasks across multiple scales (3). Dysregulation of condensate biology has been implicated in many pathological conditions, including neurodegenerative diseases (4, 5).

Condensate formation is a coupling between two distinct phase transitions – density transition and percolation transition (6-8). Biopolymers involved in such processes often have an architecture of multivalent associative heteropolymers. Such polymers are commonly modelled with a “sticker-spacer” framework (9, 10). A “sticker” is a cohesive region of the polymer sequence that may engage in inter or intra-chain interactions. Two successive stickers are interspersed by less sticky linker regions known as “spacers”. When stickers of the same type interact with each other, it is called homotypic interactions. On the contrary, heterotypic interaction refers to the interactions between different sticker types. Linear multi-domain proteins connected by flexible linkers (poly-SH3, for example) serves as a prototypical example of sticker-spacer polymer where the structured domains act as stickers and the linkers behave as spacers (11, 12). Intrinsically disordered proteins (IDPs) form another important class where certain amino acids (polar or aromatic side chains) may serve as stickers while rest of the amino acids function as spacers (13). Stickers form transient physical crosslinks (“bonds”) to generate multi-chain network (“cluster”) and beyond a concentration threshold, the system undergoes a network transition known as percolation or gelation (14) when the system shows tendency to create large (infinite in thermodynamic limit) networks. With the presence of flexible spacers, such percolation is accompanied by a density transition where the percolated clusters separate out from the solution to create a “polymer-rich” dense phase.

Physics of biomolecular phase separation immensely benefits from classical theories of homopolymers (15, 16) which describes the interplay of entropy and energy in determining the mixed or demixed configuration of such systems. Classical theory predicts two outcomes: a dispersed state at lower concentrations and a single large droplet coexisting with dilute phase at higher concentrations. However, till now, the majority of in-vitro and cellular experiments revealed a multi-droplet state where the droplets dynamically exchange components with each other but rarely coalesce to become one. Multiple physical mechanisms have been proposed to explain this conundrum. Living cells operate far from the thermodynamic equilibrium by consuming energy. Such active processes may play important roles in determining the condensate size distribution (17-19). Also, certain proteins may act as surfactant (20-22) by selectively getting adsorbed to the condensate surface. Such spatial architecture stabilizes multiple condensates.

However, in-vitro systems devoid of such complexities also demonstrate long-living droplets. We have recently proposed that such droplets exist in a dynamically arrested metastable state (8) caused by saturation of sticker valencies. The interplay of diffusion and intra-cluster rearrangement timescales determines the size distribution of condensates. The concept of dynamic arrest has been invoked recently in explaining observations regarding multi-layered condensates (23) or assembly of membrane-bound condensates (24). Competition between nucleation and coalescence kinetics (25) is also proposed as a mechanism to regulate the condensate size distribution. Additionally, multiple computational studies (26-29) highlighted the effects of inter-sticker dissociation kinetics on condensate properties like diffusivity and viscosity.

Despite all the existing efforts, a generic mechanistic understanding underlying the coalescence of condensates is incomplete. In the current study, we seek to reveal such mechanisms. What is the physical force that drives the fusion of two droplets? What is the role of sticker-spacer architecture in steering such processes? How similar are these drivers compared to homopolymers?

Using a heterotypic system, composed of two types of sticker-spacer model polymers, we have identified two distinct mechanisms underlying the kinetic arrests of condensates. Here we report that the relative energetic contributions of stickers and spacers is instrumental in determining “fusibility” of condensates. We show that fusion is governed by an intricate interplay of energy and entropy, and the sticker saturation effect triggers a kinetically arrested metastable state that prevents two condensate droplets from fusing. In contrast, homopolymers undergo a density-mediated arrest where two tightly packed condensates cannot inter-penetrate. All in all, this study provides a comprehensive mechanistic picture of physical factors that determine fusion and dynamic arrest of condensate droplets in biopolymers undergoing liquid-liquid phase separation.

## Materials and methods

### 1. Model Construction

#### 1.1 Model components

We used bead-spring polymers for coarse-grained representation of biomacromolecules (proteins and nucleic acids), with similar force-fields as our previous studies (8, 30). We considered a pair of sticker-spacer polymers (Fig. 1A) where stickers engage in heterotypic interactions (red + cyan), but homotypic interactions (red + red or cyan + cyan) do not lead to bonds between stickers.. Each polymeric chain contains 35 beads (5 stickers + 30 spacers), connected by harmonic bonds.

**Figure 1:**
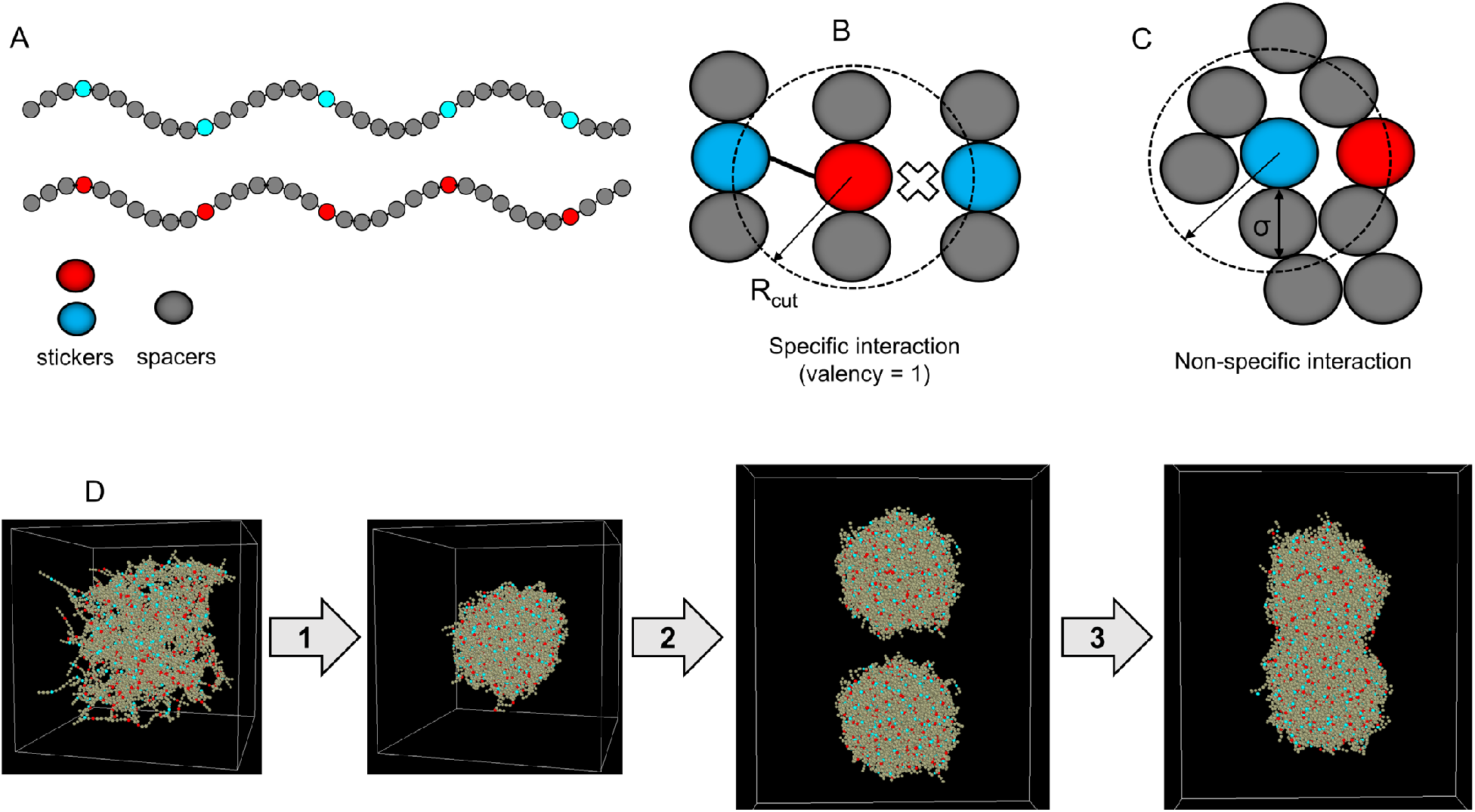
Workflow of the two-cluster fusion simulation by metadynamics approach. **(A)** Illustration of the two-component system. Each component is an associative heteropolymer consisting of 5 stickers (red and cyan beads) and 30 spacers (grey beads). We only allow heterotypic interactions, that is, red stickers interact with cyan stickers, but red-red or cyan-cyan are not allowed. **(B)** Stickers engage in specific interactions. A complementary sticker (red and cyan) pair can form a reversible bond; once bonded, they cannot engage with another sticker that may be present within Rcut. In other words, each sticker has a valency of 1. Specific interactions mimic cognate biomolecular interactions. Magnitude of specific interaction is prescribed by Es which is the depth of the harmonic well, as detailed in the method. **(C)** Spacers interact via non-specific interactions, modelled by Lennard-Jones (LJ) potential. Each bead (both stickers and spacers) can exert a long-range attractive force (within a cut-off radius, dotted circle) and short-range repulsive force which determines the bead diameter (σ). One bead can interact with multiple beads, permitted by volume exclusions. The depth of the LJ well (detailed in method) is termed as Ens which determines the magnitude of non-specific interactions. **(D)** Setup of the well-tempered metadynamics simulations. In step 1, we start with 200 uniformly distributed chains (100 red-types + 100 cyan-types) and bias the system, along the order parameter - Rg_system_, to condense into one large cluster. Rg_system_ represents the radius of gyration of the entire system. In step 2, using standard Langevin dynamics (without any bias), we relax the cluster with a pair of Es and Ens. The relaxed cluster is then copied with an initial separation of 3*Rg_cluster_, where Rg_cluster_ is the radius of gyration of the relaxed configuration. Finally, in step 3, we make the two clusters fuse with a biasing potential along the distance between cluster centers. Step 3 uses same energy pair (Es, Ens) as in step 2.

Since the focus of the current study is to derive generic principles underlying condensate coalescence, we did not attempt to model any specific system. However, the engineered protein systems like SH3-PRM or SUMO-SIM are best examples of such heterotypic systems (31) where stickers of complementary types form cross-links to create intra-condensate networks. Since the aforementioned model systems inspire our study, we limit our scope to purely heterotypic interactions. As most biological condensates comprise of multiple components (32), our two-component heterotypic system serves as a minimal model which can capture the essential physics underlying the multi-component condensates. We note that lessons learned from these simulations will readily be applicable to homotypic (single component) condensates as long as the biopolymer in question conforms to a sticker-spacer type of architecture.

#### 1.2 Polymer force-Fields

To ensure connectivity within a chain, intra-chain beads are connected by harmonic springs. Stretching energy of each harmonic bond, *E*_*bond*_ = *K*_*b*_ *(*R*−*R*_0)_^2^ where *K*_*b*_ is the spring constant and *R*_0_ is the equilibrium bond distance. R measures the distance between the bonded beads at any given time. In our model, *R*_0_ = 10 Å and 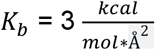.

To allow chain flexibility, angle (*θ*) between three successive beads is modelled with a function: *E*_*bending*_= κ*(1−*cosθ*)where κdetermines the bending stiffness. In our model, κ= 2 *kcal*.*mol*^−1^.

#### 1.3 Modeling specific interactions and detailed balance

To encode “specific” interactions between complementary sticker types, we introduced reversible bonds (Fig. 1B). When two stickers approach each other within a cut-off radius (R_cut_, Fig. 1B), they can form a “bond” with a probability, p_on_. The bond can be broken with a probability of p_off_, if the distance, *R*≥*R*_*cut*_. These are “specific” saturating interactions because once a sticker pair is bonded, they can’t form another bond (Fig. 1B) with complementary stickers that are still within R_cut_. In other words, each sticker has a valency of 1. The inter-sticker bonds are modelled with a shifted harmonic potential (Fig. S1A):

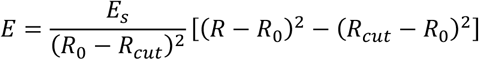

R is the inter-sticker distance. At the resting bond distance (*R*_0)_, the energy is − E_*s*_. We refer to this well depth parameter as <u>specific energy</u>. When two complementary stickers form a bond, the gain in energy is Es. In other words, the depth of energy potential is Es at the resting distance. We also note that, at *R*= *R*_*cut*_, E= 0. For *R*>*R*_*cut*_, E is set to be zero. In our model, *R*_0_ = 1.122 * σ, σ = 10 Å,*R*_*cut*_ = R_0_ + 1.5Å, *p*_*on*_ = 1, *p*_*off*_ = 1.

Since both probabilities (p_on,_ p_off_) are set to 1, the stochasticity of inter-sticker binding and unbinding is absent. For a probability < 1, there is a stochastic factor that determines whether to make or break the bond even when the distance criteria is satisfied. In our case, the bond formation or breakage only depends on the inter-sticker distance. The lifetime of the bond becomes a function of Es, such that, 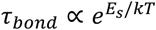. In our simulation protocol, we update (break or form) the bond statistics once in every 20 timesteps. This ensures that the newly formed inter-sticker bonds get enough time to converge to a relaxed configuration. Indeed, the inter-sticker dissociation events decay exponentially with higher Es (Fig. S2), consistent with an Arrhenius-like rate expression, *Rate* ∝ *e*^−*Es*/*k*T^. This also indicates that lifetime of individual bonds is sufficient to ensure thermalization within the harmonic well (whose depth is Es) such that detailed balance is obeyed. Since the inter-sticker association rate is a number (diffusion-limited process) determined by the particle diffusions and dissociation is a process that requires overcoming the energy barrier of the sticker-sticker bond well, the Arrhenius rates (Fig. S2) are indicative that stickers are thermalized in their corresponding wells.

#### 1.4 Modeling non-specific interactions

Apart from inter-sticker interaction, each pair of beads (stickers + spacers) interacts via a non-bonded isotropic interaction (Fig. 1C), modelled by Lennard-Jones (LJ) potential:

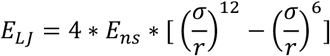

where σ represents the bead diameter and r is the separation between the beads. E_ns_ is the depth of LJ energy-well (Fig. S1B) that determines the strength of attractive potential. To distinguish it from specific interaction (described above), we will refer to this parameter as <u>non-specific energy</u>. The LJ potential enforces short-range repulsion (excluded volume) as well as long-range attraction. To achieve computational efficiency, the LJ potential is truncated at a cut-off distance (*R*_*max*_). In our model, σσ = 10 Å,*R*_*max*_ = 2.5 * σ.

It is important to emphasize the qualitative difference between specific and non-specific interactions. Unlike specific interactions, beads governed by LJ potential can form multiple “contacts” with neighboring beads permitted by volume exclusions (Fig. 1C). The magnitude of Ens (depth of the LJ well) decides the dwell time, that is, how long a group of interacting beads spend time on each other’s vicinity. On the other hand, any two stickers can have one bond at a time and the magnitude of Es (depth of harmonic wells) controls the bond lifetime. When unbonded, stickers are influenced by Ens in the same way as spacers. When they form a bond, Ens gets turned off and overridden by Es. Upon breakage of the bond, Ens again becomes operative. In this paper, we will use the nomenclature “bonds” and “contacts” to refer to the specific and non-specific interactions, respectively.

#### 1.5 Unit of interaction energies

In our simulation, we specify the interaction energy (Es and Ens) in the unit of kcal/mol. While reporting them, we express the energies in the unit of thermal energy or k_B_T where k_B_ is the Boltzmann constant, and T is the absolute temperature of the system. We will, hereafter, simply use “kT” for notational simplicity. We know that 1 kT ∼ 0.6 kcal/mol.

### 2. Simulation protocols and metadynamics

We have used the LAMMPS package (33, 34) to perform Langevin dynamics simulations of our two-component polymer system. Langevin dynamics captures the Brownian motion of particles by introducing a stochastic force at each timestep, on top of standard Newtonian dynamics. We performed our simulations in a cubic box (fixed volume) with periodic boundary conditions. Simulation temperature is 310 Kelvin. Mass of each bead is set to 1000 Dalton which roughly corresponds to 10 amino acids. With this spatial resolution, a chain composed of 35 beads refers to 350 amino acids. The viscosity of the simulation medium is described with a “damp” parameter which is set to 500 femtoseconds (fs). The “damp” parameter is inversely proportional to the viscosity of the solvent. Simulation timestep = 30 fs.

We have used well-tempered metadynamics simulations (35, 36) to facilitate the processes of polymer coalescence and cluster fusion (Fig. 1D). Metadynamics is an enhanced sampling scheme where the auxiliary gaussian potentials are imposed along a user defined order parameter (also known as collective variables, reaction coordinates etc.) to reconstruct the free energy profile of the process of interest. We have previously employed metadynamics (30, 37) to study how proteins self-assemble into liquid or solid-like condensates. In this work, we performed metadynamics simulations with two different order parameters (Fig. 1D). First, we used 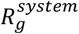 as the order parameter to bias (100 million timesteps, Fig. S3A) the coalescence of 200 uniformly distributed chains into one large cluster (step 1, Fig. 1D). 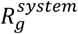 is the radius of gyration of the center of masses of 200 chains. To be precise, we considered the 17^th^ (middle) bead of each chain as its center and used their locations to compute the 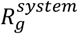. For this step, Es = 10kT and Ens = 0.5kT. The fully clustered state represents the free energy minimum (Fig. S3B). We then used standard Brownian dynamics (without any bias, 200 million timesteps) to relax the cluster with a selected pair of energy parameters, Es and Ens. We then copied the relaxed cluster and placed the second one (step 2, Fig. 1D) at an initial separation of 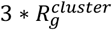 where 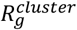 is the radius of gyration of the relaxed configuration. Using metadynamics, we then performed a two-cluster fusion simulation (step 3, Fig. 1D) with a biasing potential along the distance between the cluster centers. The center of a cluster is again defined with the location of the middle bead of each chain.

To perform well-tempered metadynamics (36), an adaptive bias is typically employed where the height of the biasing potential varies in a history dependent manner. We have used the “COLVARS” modules as a LAMMPS fix to achieve this task. We have provided <u>simulation scripts</u> that are used to setup and execute the Langevin dynamics.

### 3. Data analysis

For metadynamics simulations, we analyzed the time evolution of order parameters and related potential mean force (PMF) profile. To analyze physical properties of the clusters (density, sticker saturation etc. in Figure 3), we used the configuration files (“restart” files in LAMMPS) containing information of coordinates and topology of the system. The topology information is converted into a network. Average properties like cluster density and degree distribution of nodes (sticker saturation) are then extracted from the network.

To calculate the bond exchange entropy (discussed in Fig. 5C), we used the distribution of bonds inside (intra) and between (inter) the clusters. We divide the total bonds into three states or “labels” – cluster11, cluster22 and cluster12, where 1 and 2 are the cluster indices. So, each bond may belong to one of the three states. The bond distribution of the system is then characterized by an information entropy, *H* = −[p_11_*log*_11_+ *p*_22_ *log*_22_ + *p*_12_ *log*_12_], where *p*_11_, *p*_22_, *p*_12_ compute the probability of a bond being intracluster1, intracluster2 and intercluster respectively. For this three-state representation, p = 0.33 for a fully mixed (fused) configuration; hence maximum entropy, H_max_ = 1.1.

To compute the surface-to-volume ratio (discussed in Fig. 5F), we considered the coordinates of all 14000 beads (400 chains) and fitted it to a convex hull. We then computed the surface and volume of the hull at multiple timepoints to get the time course.

### 4. Software

#### Moltemplate

To create the model polymers, we made use of the Moltemplate (38) package which enables the user to create multiple types of chains in a template based manner.

#### Packmol

The polymers are packed inside the simulation volume using the PACKMOL (39) package.

#### LAMMPS

The Langevin dynamics simulations are performed using the LAMMPS software package (33). We used the “colvars” fix (40) to perform metadynamics, “bond/create/random” and “bond/break” fixes (41) to define the reversible bond formation within LAMMPS.

#### OVITO

We used the OVITO (basic version) software to visualize the particle motions (42).

#### Python

We used custom python scripts to setup simulations and analyze the data.

### Code availability

We have organized and released example simulations and analysis code in a public <u>github repository</u>.

## Results

### 1. Strong sticker interactions yield experimentally observed metastable condensates

Using a series of two-cluster fusion simulations, we first seek to understand how the fusion tendency of clusters depends on strength of sticker interactions (Es). The fusion dynamics is observed with a reduced parameter – “relative distance” or *R*_dist_ (Fig. 2A).

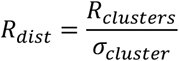

*R*_*clusters*_ is the inter-cluster distance normalized by the cluster diameter (σ_*clusters*_). When *R*_dist_ ∼ 1, two clusters are in contact, and it gradually goes down as the fusion proceeds. For a completely fused state, *R*_dist_ = 0. We note that the analysis depends on the sphericity of the clusters. For perfectly spherical clusters, R_dist_ = 1 reflects the surface contact. Our simulated clusters are far from perfect spheres, so R_dist_ provides an approximate measure of the extent of cluster penetration.

**Figure 2:**
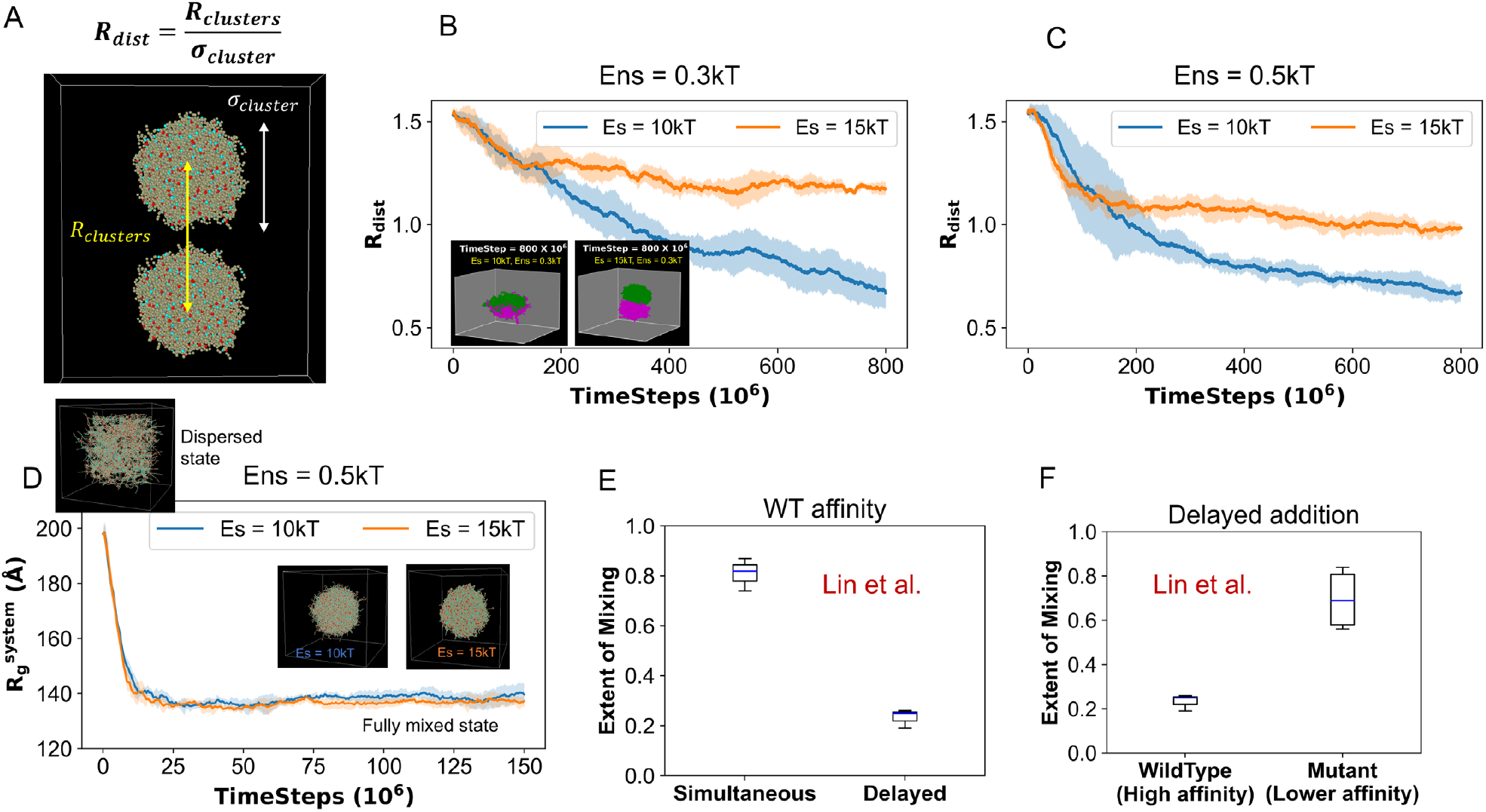
Simulation informed metastability correlates with experimental observation. **(A)** We express the inter-cluster distance with a dimensionless “relative distance” parameter: *R*_*cluster*_ which is the inter-cluster distance (*R*_*cluster*_) normalized by the cluster diameter (σ_*cluster*_). **(B)** Fusion behavior of clusters under stronger (Es = 15kT, orange line) and weaker (Es = 10kT, blue line) specific interaction strengths. Non-specific interaction strength, Ens = 0.3kT. Each trajectory is an average over 5 stochastic runs (Solid line: mean, fluctuation envelop: standard deviation). Insets display the representative snapshots of last simulation timeframe (parameters mentioned in the labels). Two clusters are colored differently for visual clarity. **(C)** Same configurations as B, except Ens = 0.5kT. **(D)** Clustering dynamics of 400 uniformly distributed chains into one large cluster, at two parameter combinations (Es = 10kT and 15kT with Ens = 0.5kT). Insets show snapshots depicting dispersed (Initial state, upper left) and fully-mixed or fully-clustered (lower right)states. Each line is an average of 5 stochastic trials. **(E, F)** Experimental data replotted from (41). The three-component experimental system (detailed in texts) contains two types of RNAs (BNI1 and CLN3) and a protein Whi3. In E, authors first mixed all the three components at the same time (labelled as “simultaneous”) and measure the colocalization of the RNAs. Then they mixed Whi3 + BNI1, wait for 4 hours, and then added CLN3 (labelled as “delayed”). The colocalization of two RNA types are quantified with Pearson’s *r*-values, which we have plotted on the vertical axis as “Extent of Mixing”. To display the distribution, we used standard boxplot representation or “five-number summary” consisting of the minimum, the maximum, the sample median, and the first and third quartiles. In **F**, only the delayed protocol is shown. Here, the authors used a mutant BNI1 which has a reduced affinity for Whi3. So, the wildtype is Whi3 + BNI1 and then CLN3, while the mutant version is Whi3 + BNI1_mutant and then CLN3.

Firstly, we delineate the phase transition boundary from the cluster relaxation dynamics (Fig. 1D, step 2). When we relax (without any bias) the cluster under different pairs of specific and non-specific energy parameters (Es and Ens), the minimum combination (critical level) that yields a stable cluster is Es = 8kT and Ens = 0.3kT (Fig. S4). For any pair of Es and Ens below that critical level, the cluster falls apart or dissolves. Thus, above the critical level, we have selected four combinations of Es and Ens (Es = 10kT, 15kT; Ens = 0.3kT, 0.5kT) to explore how cluster fusion varies at weaker and stronger values of Es and Ens.

Fig. 2B shows that, with stronger Es (15kT), two clusters do not fuse (orange line, *R*_dist_ > 1) even though they touch each other (Inset panel). At a relatively lower Es (10kT, blue line), clusters merge readily. The Movie S1 demonstrates these distinct merging patterns where the green cluster keeps mixing with the magenta one at Es = 10kT but fails to merge at Es = 15kT (arrested state). We notice that the 10kT fusion profile (blue line, Fig. 2B) tends to go down even at the last timepoint, which means we need to wait longer to observe the completely fused (R_dist_ = 0) state. The 15kT line, on the other end, fluctuates above the R_dist_ = 1 level for a long time. These behaviors are replicated at a different non-specific interaction strength, Ens = 0.5kT (Fig. 2C). From Fig. S4B, it is worthy to note that Ens = 0.5kT can stabilize a cluster even at Es = 0, indicating a pure non-specific energy driven condensation phase transition. With this context, it is more interesting that stronger Es can cause an arrest even for clusters that are stable without sticker-sticker bonds. Despite qualitative similarity, there are subtle differences between the fusion profiles. For Ens = 0.5kT (Fig. 2C), the Es = 15kT line runs closer to the Rdist = 1 level, compared to the Ens = 0.3kT case (Fig. 2B). These four scenarios already point towards an interplay of Es and Ens in controlling the fusion which will be later explored in greater detail.

Next, we start our simulations with 400 chains uniformly distributed in the simulation volume at two different Es (10kT and 15kT). In both cases, 400 chains coalesce into one large cluster (Fig. 2D), with a free energy minimum around the fully clustered state (Fig. S5). Under the same parameter combination (Es = 15kT, Ens = 0.5kT), two clusters, each carrying 200 chains, encounter an arrested state (Fig. 2C) while randomly distributed 400 chains can bypass such a configuration to form one large cluster (Fig. 2D). So, the initial condition of clustering events determines the outcome – one single cluster or two separate clusters that do not merge on the timescale of simulation. This is a hallmark of a non-equilibrium metastable state.

Recently, Lin et al. (32) provided an experimental evidence of metastability in biological condensates. They considered a ternary system consisting of two RNA types (RNA1 and RNA2) and an RNA-binding protein, Whi3. These heterotypic condensates showed distinct spatial arrangements based on the timing of the addition of the components. In the first set of experiments, authors mixed all three components at the same time. This “simultaneous” addition created a well-mixed condensate (Fig. 2E) where mixing is quantified by the colocalization of RNA1 and RNA2. In an alternative scheme, the authors first mixed Whi3 + RNA1, waited for 4 hours and then added RNA2. Surprisingly, this “delayed” addition yielded a significantly unmixed state (Fig. 2E) where RNA1 and RNA2 each localized in their respective homotypic condensates but did not colocalize with each other. This experimental observation completely recapitulates our model predicted metastability (Fig. 2C and Fig. 2D) where the timing of events steers the system into different configurations. To consolidate our model predictions, we compare them with another key experimental observation. Lin et al. (32) also generated a mutant version of RNA1 (BNI1_mutant) that weakly binds to Whi3. Using the delayed-addition protocol, when they repeated the assay, that is, Whi3 + mutant RNA1 (low affinity), wait for 4 hours and then add WT RNA2, they observed a “rescued” mixing (Fig. 2F). The low-affinity mutant of RNA1 promoted the recruitment of RNA2 into the pre-existing condensates. This observation directly correlates with the differential fusion profiles (Figs. 2B and 2C) of clusters at two different sticker-sticker interaction strengths (Es = 10kT and 15kT). Just like the higher protein-RNA affinity causes arrest in the ternary experimental system, a stronger Es between the complementary stickers of our model proteins triggers an arrested state where two clusters can no longer merge to become one.

## 2. Kinetic arrest stems from long-living inter-sticker bonds

Next, we elucidate the physical properties of the individual chains that can be mapped into their fusion tendencies (Fig. 3). As shown in Figs. 2B and 2C, we consider four parameter (Es, Ens) combinations, labeled C1-C4 – two of them undergo fusion (C1 and C2, labelled in green) and the other two display an arrest (C3 and C4, labelled in red). Density of clusters correlates with the strength of non-specific interactions (Ens) between spacers (Fig. 3A) and does not predict the fusion outcome. In fact, clusters with similar density (C2 and C4) show very different fusion behavior. Surprisingly, despite having lower density, C3 undergoes arrest while denser C2 merges readily. When we look at the degree of sticker saturation within the clusters (Fig. 3B), a trend emerges. For C1 and C2, weaker Es results in more free stickers, while stronger Es pushes the system towards 100% sticker saturation as in the case with C3 and C4. It is interesting to note the effect of spacer energetics (Ens) in stabilizing the Es-mediated bonds. For instance, higher Ens (C1 and C2, Fig. 3B) promotes higher degree of sticker occupancy (for identical Es), by changing the density of the system. However, even though C2 and C3 have overlapping distributions, they show distinct fusion behaviors. Analyzing the connectivity within the polymer network (Fig. S6) does not yield any causal trend. When we look at the radial distributions of sticker saturation, for Ens = 0.3kT (Fig. S7A), cluster with 10kT Es has a radial gradient of sticker saturation that is absent in the 15kT case. More stickers are free towards the surface than the cluster core which might initiate the merging process with another cluster having similar configuration. This effect is again obscured when we examine the 0.5kT Ens scenario (Fig. S7B) where the difference between 10kT and 15kT is not significant.

**Figure 3:**
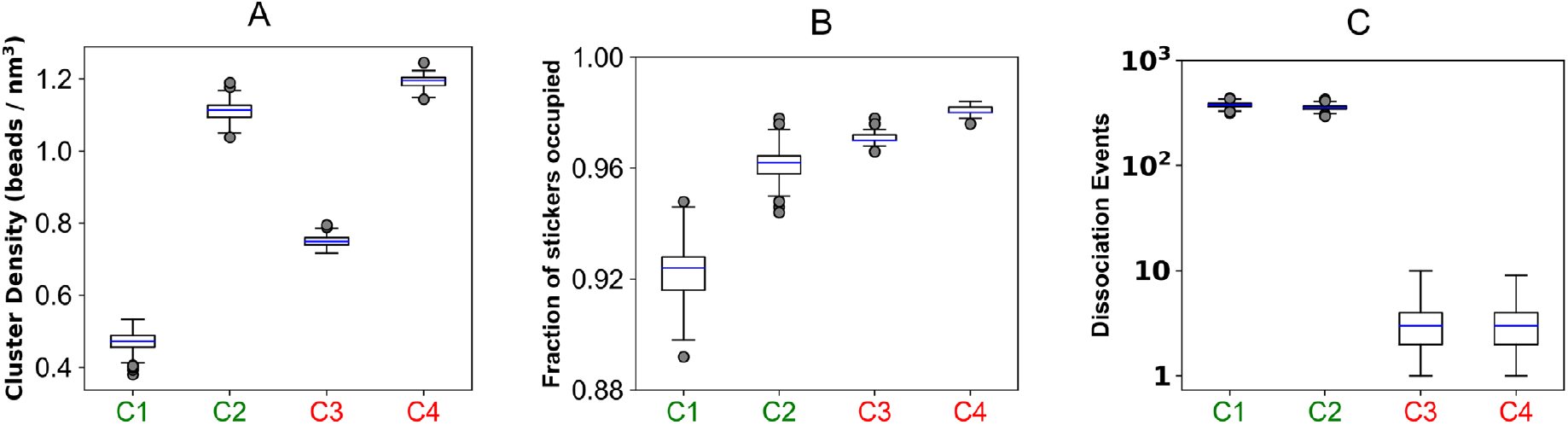
Mapping fusion behavior to physical properties of the clusters. Parameter combinations (Es, Ens) are divided into two categories: C1 (10kT, 0.3kT) and C2 (10kT, 0.5kT) are labelled in green as they undergo fusion; C3 (15kT, 0.3kT) and C4 (15kT, 0.5kT) are labelled in red which do not fuse. **(A)** Density of clusters under four parameter combinations. 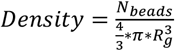 where *R*_*g*_ is radius of gyration of the cluster and *N*_*beads*_ is the total number of beads (stickers + spacers) present in the cluster. **(B)** Fractional occupancy of stickers that indicates, on average, what fraction of the total sticker population are bonded. **(C)** Dissociation events refer to how many times an inter-sticker bond is broken in between two successive observation points. Events are plotted in log scale. To derive the distributions (A, B, C), for each condition, we run 10 stochastic trials and sample 10 timepoints from each trial; hence each distribution is collected over 100 snapshots. To display the distribution, we used standard boxplot representation or “five-number summary” consisting of the minimum, the maximum, the sample median, and the first and third quartiles.

The most striking correlation we observe when we quantify the dissociation rates of stickers (Fig. 3C). This rate reflects the extent of bond re-organization within the cluster. Since breaking of inter-sticker bonds involves overcoming the specific energy well (whose depth is Es), number of bond dissociation decays exponentially with higher Es, as highlighted in Fig. S2. This slower dissociation rate affects the internal dynamicity of the cluster.

To probe the effect of dissociation kinetics, we gradually increase the sticker content of the chains (Fig. S8A). The number of stickers per chain is termed as valency. Keeping the interaction energies identical (Es = 10kT, Ens = 0.3kT), as we increase the valency, we see a gradual upward shift in cluster density (Fig. S8B) and sticker saturation (Fig. S8C). Interestingly, dissociation events per sticker reaches a plateau (Fig. S8D), instead of declining gradually. As a result, clusters made of higher valent chains still undergo fusion (Fig. S8E).

Comparing Figs. 3A-3C, along with Figs. S6, S7, S8 and S9, we notice that static properties of clusters (density, network architecture, sticker occupation) cannot predict the arrest tendency. It is rather a kinetic effect that originates from the lifetime of a bond, that is, how fast a bond can break and reform. Thus, we conclude that strong specific interactions trigger a kinetic arrest where the stickers form a saturated network with *long-lived bonds*.

### 3. Separation of sticker-spacer energetics yields re-entrant merging behavior

Next, we explore the interplay of sticker and spacer energetics in determining the cluster fusion behavior. Specifically, we seek to explore whether there is any difference in fusion of clusters composed of heteropolymers (sticker-spacer) vs homopolymers (spacer only). By tuning the gap between Es and Ens, we can systematically create a spectrum of polymer systems where the energy separation between stickers and spacers are gradually altered.

Fig. 4A displays the phase diagram where extent of fusion is computed against a wider combination of sticker spacer energetics. It is important to note that the diagram reflects a “kinetic phase space” which will slowly change over time since we are dealing with kinetically arrested states. But for a pragmatic purpose, we simulate each system for a given time (which is long enough to demonstrate slowed fusion) and report the phase at the last point.

**Figure 4:**
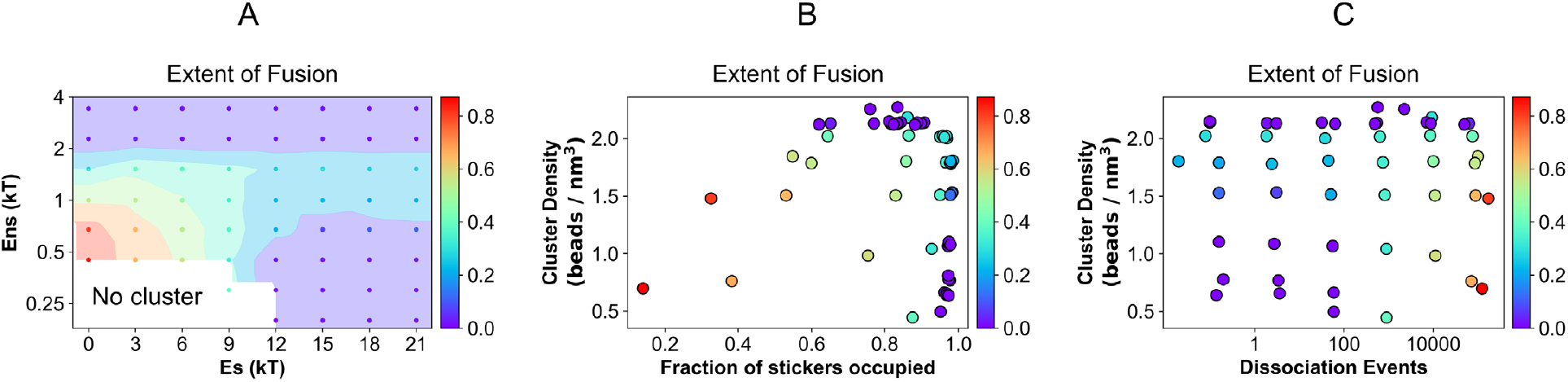
Phase diagram of inter-cluster fusion. **(A)** Fusion of clusters as a function of a wide range of specific (Es) and non-specific (Ens) interactions. The colored circles are the points where the simulations are carried out. These discrete points are then extrapolated to generate the continuous phase space encoded by the color scheme. At the lower left region, clusters are unstable. The fusion tendencies are mapped to the two-dimensional plane of **(B)** cluster density vs extent of intra-cluster sticker saturation, and **(C)** cluster density vs bond reorganization rates. The color bar displays the extent of fusion which is defined as 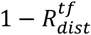, where 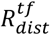 is the relative inter-cluster distance at the final timeframe of the simulation. Each point is an average over 5 stochastic trials, where each trial is executed for 400 million timesteps.

First, we notice that the lower left corner of the diagram has no cluster (density is displayed in Fig. S10), signifying a zone below the phase transition limit. The leftmost column (Es = 0) represents clusters made of homopolymers. Just above the “no cluster” zone, cluster fusibility is the highest as shown by the red region. For Es = 0, as we titrate up Ens, merging tendency gradually goes down and an arrest is encountered at Ens ≥ 2kT. Similarly, if we consider the row at Ens = 0.45kT, fusion propensity is very high at Es = 0. Moving along the higher Es gradually decreases the fusibility and the system enters an arrested state at Es ≥ 12kT. Sticker driven clusters (high Es, low Ens) occupy the lower right zone of the diagram which belong to the arrested phase entirely. If we move up from this zone, we find a very interesting non-monotonic trend. Kinetic arrest is partially rescued at an intermediate Ens. But clusters re-enter the arrested state at larger Ens.

When we map this fusion behavior into the sticker saturation vs. cluster density space, the trend becomes clearer (Fig. 4B). The arrested state (purple points) appears either on high density zone or greater extent of sticker saturation region. Looking at the plane of density vs sticker dissociation rates (Fig. 4C), apart from the high-density region, purple points lie at the lower dissociation zone. Combining Figs. 4B and 4C, we conclude that there are two distinct mechanisms of kinetic arrest at play here. The density mediated kinetic arrest for homopolymers is a fundamentally different mechanism than the sticker saturation mediated kinetic arrest for sticker-spacer polymers. It is noteworthy that density causes arrest for sticker-spacer polymers even when the inter-sticker bonds remain highly dynamic (upper-right purple points in Fig. 4C).The interplay of these two mechanisms (density vs sticker-saturation) yields the re-entrant fusion behavior which can be better explained once we uncover the driving forces of the cluster merging in the next section.

### 4. Interplay of energy and entropy drives the fusion of clusters

We have shown that the loss of dynamic bond organization prevents fusion of two clusters composed of sticker-spacer polymers. For homopolymers, it is the cluster density that triggers the arrest. But what is the physical force that drives the fusion of clusters? To answer that question, we consider fusion trajectories for a heteropolymer (Es = 10kT, Ens = 0.3kT) and a homopolymer (Es = 0kT, Ens = 1kT) system (Fig. 5). For heteropolymers, firstly we note that the potential energy does not change much as the fusion proceeds (Fig.5A). There is a slight downward trend in potential energy that mainly comes from the non-specific interactions (E_contact_ in Fig. S11A) between spacers. It is important to recall that at Es = 10kT and Ens = 0.3kT, cluster is stabilized by sticker-sticker interactions. Strikingly, we notice that the total number of inter-sticker bonds in the system does not change during the course of fusion (Fig. 5B); the number of inter-cluster bonds gradually rises at the expense of intra-cluster bonds. The quantitative effect of this bond exchange can be captured by an information entropy, which is called bond exchange entropy hereafter (Fig. 5C). This parameter describes how the total number of bonds are distributed across the two-cluster system. As the clusters interchange bonds, exchange entropy keeps rising. For a fully fused state, entropy has a maximum value of 1.1 (described in Methods). Since the change in potential energy is minimal and total number of bonds remains constant, the fusion process has to be driven by entropy. Although the exchange entropy defined here does not capture all components of the thermodynamic entropy (translation, conformations etc. of polymeric chains), it persistently goes up as the fusion continues to happen. Now, when there are not enough inter-cluster bonding opportunities due to intra-cluster sticker saturation, fusion comes to a halt due to the lack of bond exchange (Fig. S12). Analysis of neighbor exchange (Fig. S13) reveals that inter-cluster neighbors, on average, exceed one as the fusion proceeds. For a case with kinetic arrest, chains from cluster-1 most likely cannot engage with more than one chain from cluster-2 which prevents them from merging.

**Figure 5:**
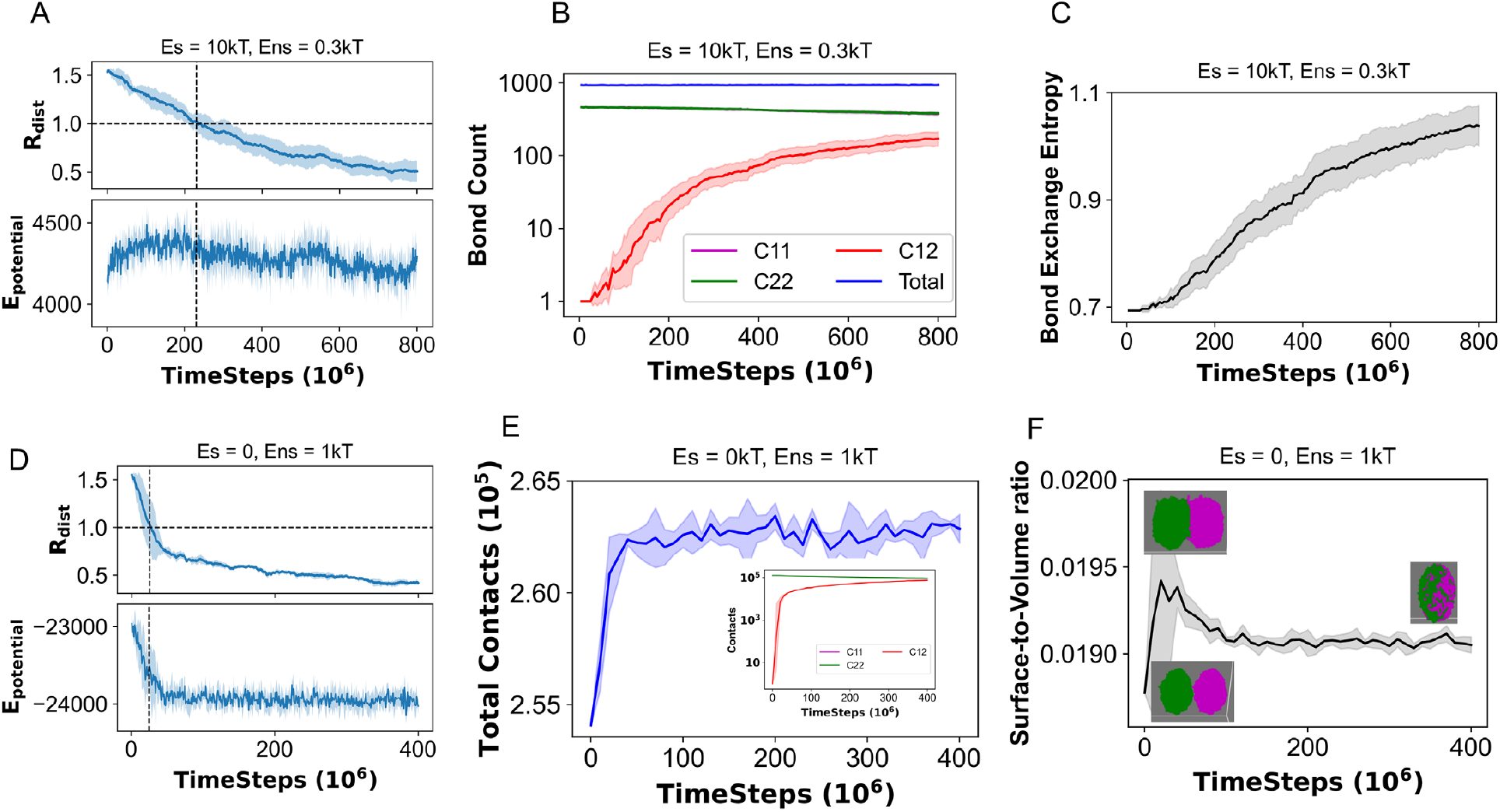
Driving forces of cluster fusion. First three panels (**A**,**B**,**C**) represent a heteropolymer system (Es = 10kT, Ens = 0.3kT), while last three panels (**D**,**E**,**F**) represent a homopolymer system (Es = 0, Ens = 1kT). (**A, D**) Potential energy profile (unit: kcal/mol) along with the cluster fusion profile. R_dist_ is defined in Fig. 2A. (**B, E**) Number of bonds and contacts, respectively. A “bond” is a cross-link between two heterotypic stickers while any two beads within a cut-off distance (22.44 Å, detailed in method) are counted to have one “contact”. C11, C22 and C12 stand for intra-cluster-1, intra-cluster-2 and inter-cluster, respectively. “Total” indicates the entire system. For **E**, contact exchange between clusters is shown as an inset. (**C**) Measure of bond exchange entropy (detailed in method) for the entire system. (**F**) Surface to volume ratio of the two-cluster system as the fusion proceeds. Each analysis is an average of 5 stochastic trials.

We now turn our attention to homopolymers. First, we see a significant drop in potential energy (Fig. 5D) as the clusters begin to fuse; however, after the initial drop, potential energy plateaus even though the fusion continues to happen (upper panel, Fig. 5D). The potential energy profile is again shaped by the pairwise contact energy (Fig. S11B). This drop in potential energy is accompanied by a gain in contact counts (Fig. 5E). When two clusters begin to merge, a fraction of the surface beads establish new contacts. Since Ens is relatively high (1kT), this increase in contacts favors an energy driven fusion. Movement of beads from surface to interior manifests as a reduction of surface tension. Fig. 5F shows the trend in surface to volume ratio as fusion starts to happen. When two clusters approach from a distance, the combined system has a cylindrical shape which explains the initial hike in the trend. Once they are in contact, inter-penetration of two clusters results in a decrease in surface to volume ratio. Movie S2 helps us to visualize the difference in fusion kinetics between a homopolymer and a sticker-spacer polymer. Due to the surface tension effect, despite having higher density, homopolymers initiate merging faster while the relatively loose clusters made of sticker-spacer chains penetrate slowly by gradual exchange of polymers.

We also notice that the potential energy, total contact counts and surface-to-volume ratio all converge to steady levels around 100 million timesteps, but the fusion parameter (R_dist_) continues to go down. This suggests an entropic contribution towards the later stage of fusion. We can, again, invoke the concept of an exchange entropy to rationalize this phenomenon. The steady rise of inter-cluster contacts (Fig. 5E, inset) indicates that the exchange of beads drives the process towards completion. Hence, we observe a combined effect of energy and entropy. The initial energy drop clearly shows an energy inceptive in initiating the fusion and the contact exchange between the clusters later serves as additional incentive to drive two clusters into a fully fused state. When Ens is very high, clusters become too dense (solid-like). In this scenario, beads from one cluster can no longer flow into the other, causing a kinetic arrest.

## Discussion

In this work, we have identified two distinct physical mechanisms that may potentially trigger kinetic arrest of biomolecular condensates. Sticker spacer heteropolymers may undergo a sticker saturation mediated arrest while spacer-only homopolymers need to cross a threshold density to encounter arrest.

We first established a direct correlation between our model predictions and experimental observations (32) in terms of the existence of metastability for sticker-spacer polymers. We showed that the sequence of simulations (Figs. 2B – 2D) yields qualitatively different results that can be mapped to the timing-dependent condensation behavior of heterotypic protein-RNA condensates (Figs. 2E, 2F). Our simulation also rationalizes the time-dependent material properties of homotypic (FUS or PGL-3) condensates (43) where the fusion tendency gradually goes down as the condensates mature or age. Due to the reorganization dynamics of stickers, there exists a time-window where a fraction of the stickers may not be saturated, and condensates can merge at this stage. With relaxed configurations, most of the stickers are in a saturated state, and these “matured” condensates can no longer merge.

Having established experimental correlations, we sought to unravel the physical driving forces underlying the merging of condensates. In this process, we uncovered an interesting interplay of sticker and spacer energetics in driving the clustering of biopolymers. Fig. 6 summarizes our mechanistic understanding. For a solution containing associative polymers, above a system-specific threshold concentration, the system will separate into dilute and dense phases. This phase transition is driven by energy, be it heteropolymers (Fig. S14) or homopolymers (Fig. S15). However, the relative energetic contribution may vary between the beads. One end of this spectrum is sticker-spacer polymer, while the other end is homopolymer.

**Figure 6:**
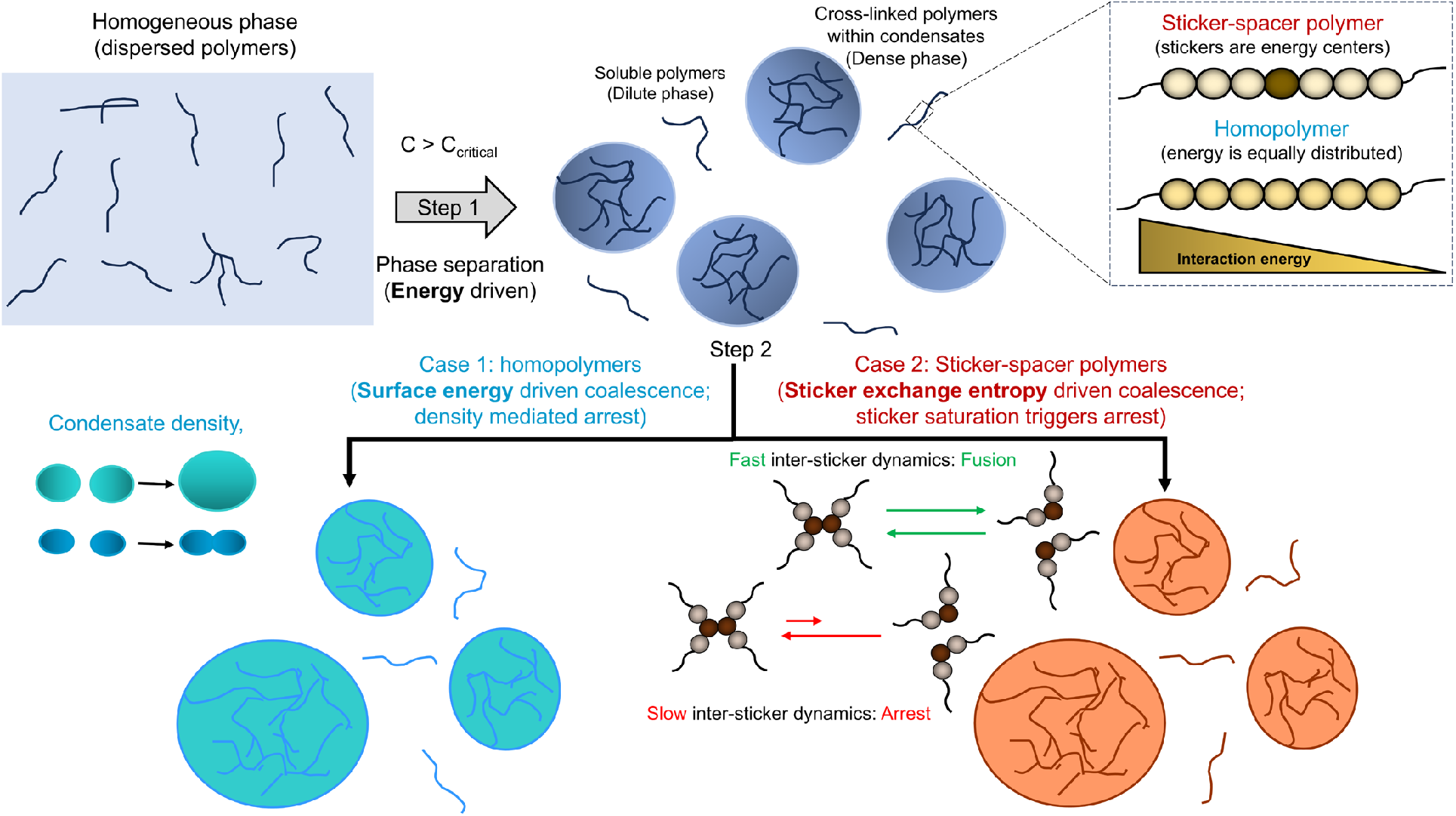
Graphical summary of two distinct physical mechanisms of kinetic arrest. Below a system-specific critical concentration, biopolymers remain in a dispersed state (monomers + oligomers) which appears as a single homogeneous phase. Upon crossing that critical concentration, the system separates into dilute and dense phase where multiple condensates emerge (**Step 1**). Now, depending on the spatial distributions of interaction energies, polymers can be classified into two categories (**inset**). For homopolymers, each bead interacts with the other beads in an equivalent manner. Hence, each bead has equal energetic contribution towards phase separation. For associative heteropolymers, stickers are the energetic drivers for phase separation. Once multiple condensates form, in the case of sticker-spacer polymers (**Step 2: Case 2**), exchange of stickers drives the fusion of two droplets. If the stickers are fully saturated (bonded with complementary sticker types) within a condensate, the entropy driven merging does not take place due to the slow bond reorganization dynamics. On the other end, two condensates made of homopolymers (**Step 2: Case 1**) merge to minimization of surface tension. This energy driven fusion comes to a halt when the density of condensates exceeds a threshold level.

For sticker-spacer polymer, sticker-sticker interactions stabilize the cluster while the flexible spacers confer liquidity and modulate density by providing weaker interactions. For such systems, we have shown that the ability of clusters to exchange stickers decides the fusion propensity. As more and more intra-cluster stickers are saturated, purely due to slowed inter-sticker dissociation, the ability of the two clusters to coalesce gradually decreases. When nearly 100% stickers are saturated or the effective cluster valency is fully exhausted, they cannot merge when come into contact. In terms of driving force, since stickers are energy centers, exchangeability of stickers are sole determinant of fusion. This scenario is akin to “*mixing of two ideal gases*” where entropy drives the gas molecules to explore more volume. The entropic gain stemming from sticker exchange drives the fusion process since most of the stickers are already in bound form and total bond count does not vary (Fig. 5B) as the fusion proceeds. Now, for a fully saturated cluster, fusion is not possible as the long-lived inter-sticker bonds do not open up readily to promote entropically driven exchange. It is important to highlight that the single valence nature of individual stickers is the causal factor underlying the kinetic arrest.

On the other hand, for homopolymers, energy is equally distributed amongst all the beads. So, exchange of each bead is equally important. At the same time, surface beads can only form partial contacts due to their peripheral locations. The radial difference in contact energy manifests in the form of surface tension. The merger of two homopolymeric clusters reduces the surface to volume ratio. In other words, the beginning of fusion transfers multiple beads from surface to interior; such surface energy minimization is a generic fusion mechanism of two liquid droplets. Since non-specific interactions are not restricted by any “single valence” effect, multiple contacts allow two clusters to exchange beads easily. Hence, for homopolymeric condensates, the surface energy initiates the merger and the entropic boost from contact exchange drives the process towards the finish line. For a large non-specific energy higher than a critical level, the density of such clusters becomes too high to flow towards each other.

The manifestation of surface tension is also different in homopolymer vs sticker-spacer polymer. A sticker can be only bonded or free, irrespective of the location (surface vs bulk of the cluster), since the valency is 1. On the other hand, spacers near the cluster surface remain “frustrated” due to less than maximum possible contacts. Upon fusion, spacers “gain” contacts, but stickers need to “exchange”. These distinct interaction modes yield an energy driven fusion for homopolymers, but exchange entropy driven merger for sticker-spacer polymers. This might also explain the order of magnitude lower surface tension of biological condensates (44), compared to the canonical oil droplets.

The full spectrum of energy scale separation explored here is crucial for biological systems. The sticker-spacer architecture in proteins and nucleic acids comes with a variety of flavors. For linear multivalent proteins, multiple folded domains (stickers) are tethered together by flexible lDRs (spacers). This architecture is starkly contrasted by intrinsically disordered proteins (IDPs) where a single amino acid residue or a collection of residues may serve as stickers connected by disordered spacers. The interaction energy between globular-globular and globular-IDR interfaces can be very different (45). This difference, in turn, may dictate the separation of sticker-spacer energetics that allows evolutionary selection of protein sequences with diverse potential functions.

Lin et al. (32) showed that the phenomenon of kinetic arrest plays important role in determining phenotype in living cells. They discovered that in vivo ablation of kinetic arrest in protein-RNA condensates causes defects in cellular morphology. This underscores the functional importance of distinct condensate compositions that are enabled by the relative energetics of stickers and spacers. Dynamical arrest also stabilizes the multi-condensate state, and the condensate size distribution can be directly tuned by the gap in energy scales. When energetic difference between stickers and spacers reduces, exchange of both stickers and spacers becomes important and sticker saturation alone can’t trigger the arrest. Conversely, stickers alone would control fusibility if they were the major energetic contributors. Our simulations also revealed the interplay between sticker-saturation and density mediated kinetic arrest that might give rise to a re-entrant phase behavior (Fig. 4A). Potentially through post-translational modifications, depending on the functional context, cellular systems may reversibly switch between arrest vs no-arrest state. Such mechanistic principles could also be relevant in creating multi-phasic or multi-layered condensates.

Work presented here highlights a clear connection between the structural details of biopolymers and the material properties of mesoscopic structures that they assemble into. Studying viscoelastic properties of condensates (46, 47) has been an active field of investigation due to its implications in biological functions. A recent study (48) showed the role of spacer mutations (glycine-to-serine) in promoting kinetic arrest as well as altered viscoelastic properties of the condensates. Our work can contextualize these observations in the light of creating sticker-spacer architecture with different degrees of energy separation, that can tune the material properties of the mesoscopic condensates.

We have explored the effects of interactions strengths for a given arrangement of stickers and spacers in a sequence. However, the patterning of stickers itself (13) could dictate the material properties and fusibility of condensates. How dynamical control might be achieved via optimized sticker patterning remains to be an intriguing evolutionary question that can be addressed in future. Another important parameter is the relative energetics of homotypic and heterotypic interactions. In this paper, we have tuned the heterotypic energy strength. Presence of homotypic interactions introduces another layer of complexity. Our previous report (8) explores the effects of non-specific interactions (Ens) on chain collapse and clustering dynamics in general. However, on the level of sticker-sticker (single-valent specific) interaction, the interplay of homotypic and heterotypic interactions may govern the competition between intra-chain and inter-chain collapses. This rich physics can be studied in future by systematically altering the homotypic vs heterotypic energy parameters.

## Author Contributions

A.C. and E.I.S. designed the research, A.C. performed simulations, A.C. and E.I.S. analyzed the data and wrote the paper, E.I.S. secured the funding.

## Acknowledgement

We are grateful to Aditya Ranganathan, Junlang Liu and Sayantan Mondal for fruitful discussions. A.C. greatly appreciates the help with LAMMPS software and metadynamics simulations provided by Aditya Ranganathan. A.C. thanks David Kanovich for providing inputs with the github documentation. This work was supported by NIH (Grant 5R35GM139571).

## Supplemental Movie Legends

**Movie S1:** Fusion dynamics of two clusters under two combinations of energy parameters. Es = 10kT shows merger while Es = 15kT undergoes arrest.

**Movie S2:** Comparison of fusion dynamics between sticker-spacer polymer (Es = 10kT, Ens = 0.3kT) and homopolymer (Es = 0, Ens = 1kT)

## Supplemental Figures

**Figure S1:**
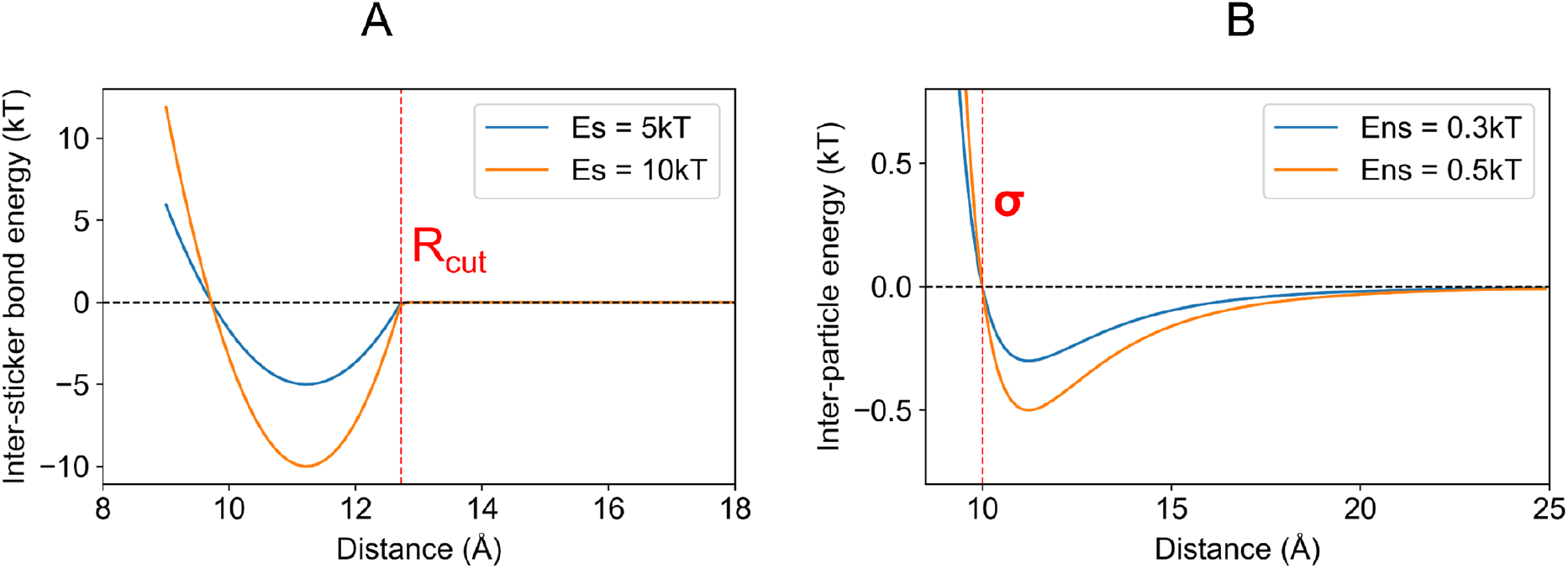
Illustration of specific and non-specific energy potentials. (A) The inter-sticker bonds are modelled with a shifted harmonic potential which becomes zero at a distance greater than R_cut_. At the resting bond distance, the gain in energy is Es (E_specific_). In other words, the depth of energy potential is Es at the resting distance. Two energy potentials are depicted for two different Es. (B) Pairwise non-specific interactions are modelled with Lennard-Jones potential which enforces excluded volume by the parameter, σ. The energy minima lie at the resting distance, that is 1.122 * σ. Depth of the energy well is Ens (E_non-specific_). Two potentials are depicted at two different Ens.

**Figure S2:**
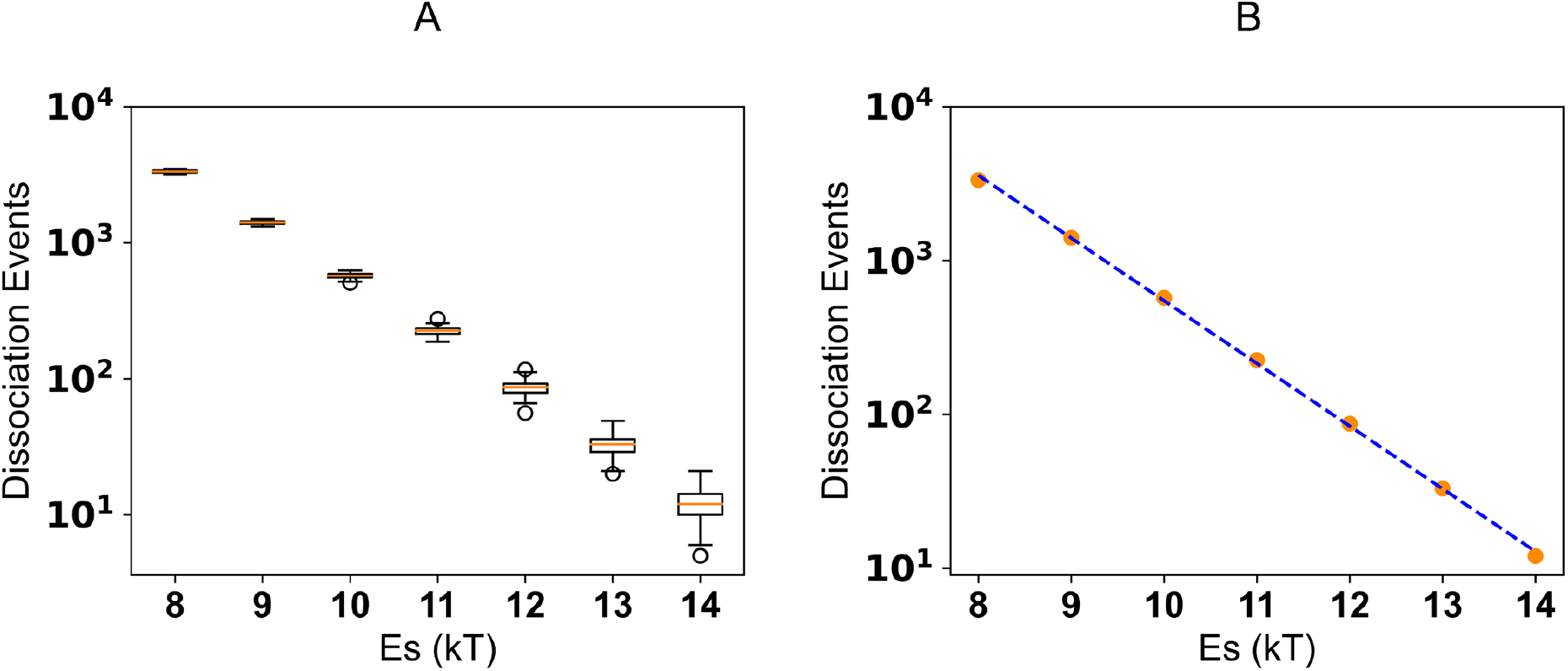
Inter-sticker dynamics follows an Arrhenius-like rate. For sticker-sticker interactions, rate of dissociation, 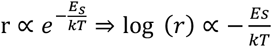. (A) The number of bond dissociation events between the red and cyan stickers (Figure 1A), at Ens = 0.3kT, as a function of specific interaction energy (Es). We note the log scale on the vertical axis. For each condition, we sampled 50 timeframes when the system is equilibrated. To display the distribution, we used standard boxplot representation or “five-number summary” consisting of the minimum, the maximum, the sample median, and the first and third quartiles. (B) The logarithm of dissociation events is fitted as a linear (blue dashed line) function of Es which yields a negative slope, consistent with an Arrhenius-like rate expression.

**Figure S3:**
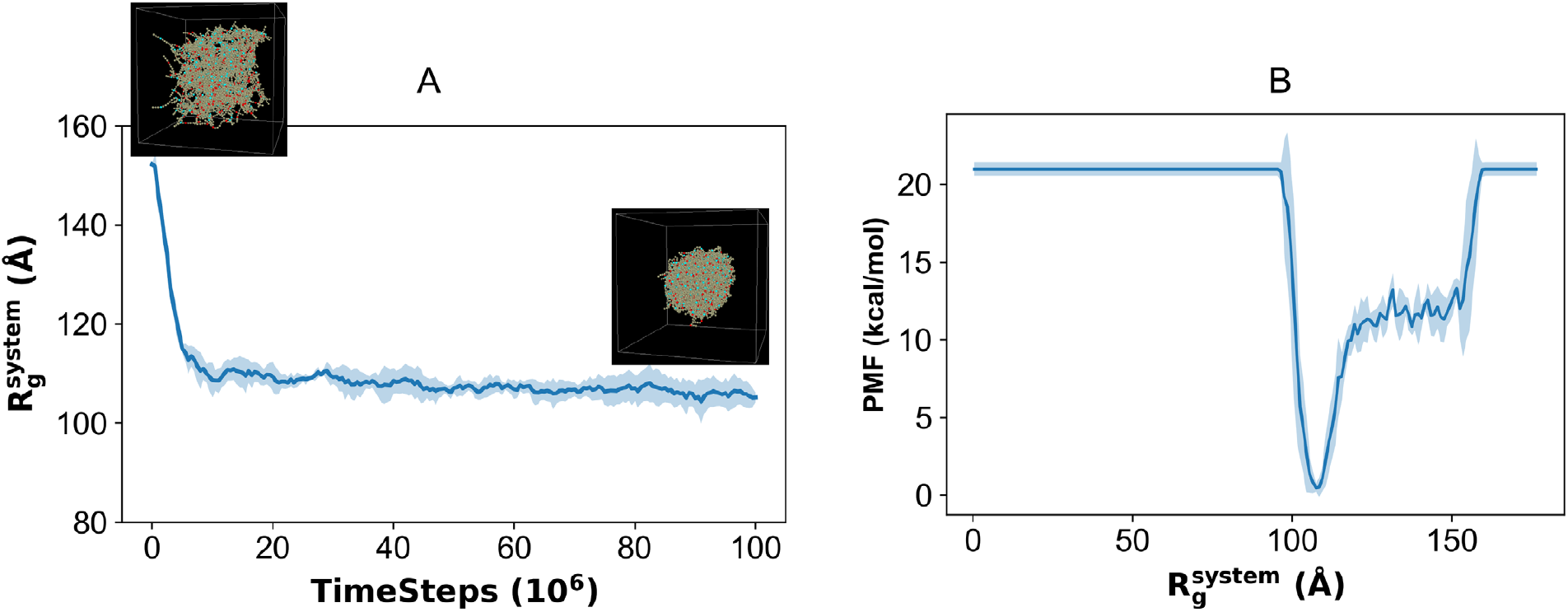
Coalescence dynamics of 200 chains into one large cluster. Energy parameters, Es = 10kT, Ens = 0.5kT. (A) Timecourse of the metadynamics order parameter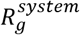. Insets show first and last timeframes depicting dispersed and clustered states. (B) Free energy profile where the minimum corresponds to the fully clustered state. Each line is an average of 5 stochastic trials. The solid line is the mean and fluctuation envelop represents the standard deviation.

**Figure S4:**
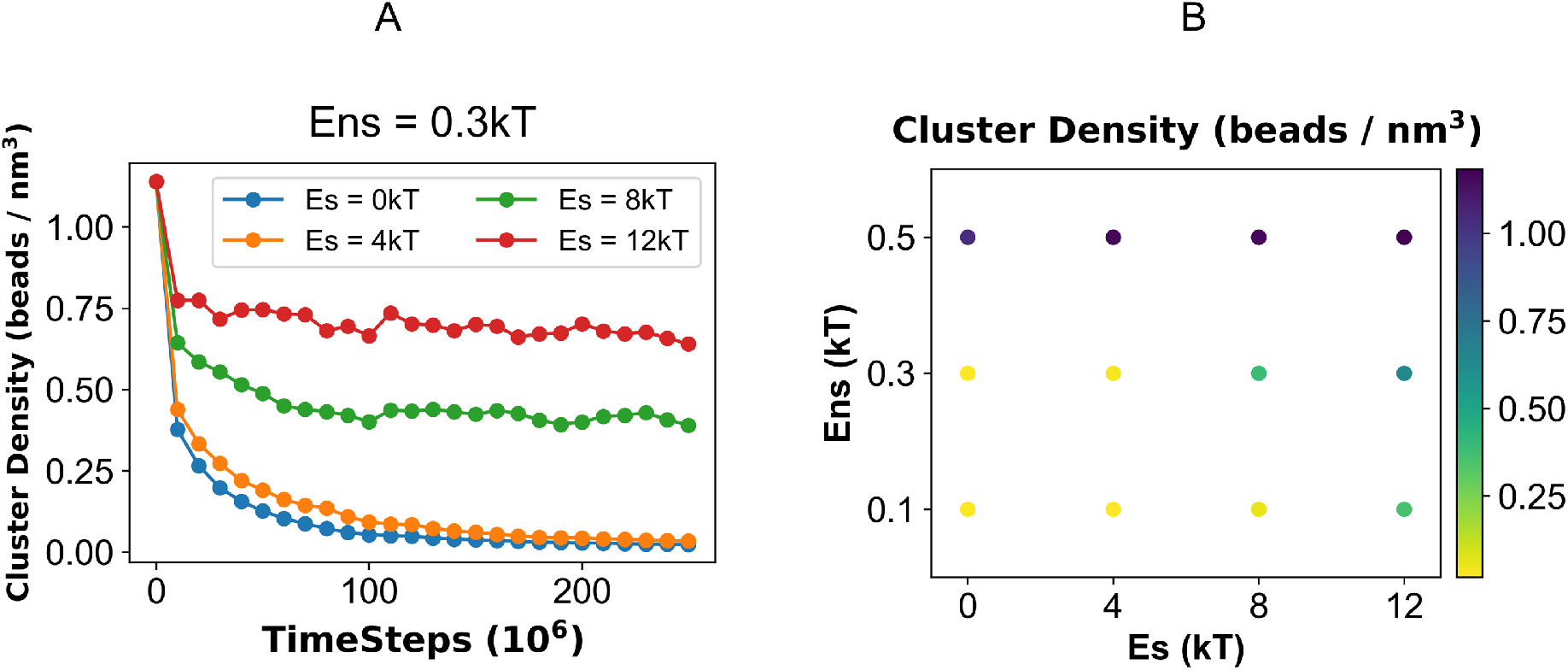
Quantification of phase transition boundary from the relaxation dynamics. (A) Timecourse of the cluster density, for a fixed Ens = 0.3kT, as a function of specific interaction strength, Es. (B) Phase diagram of the cluster density which is computed at the last timepoint of relaxation trajectory. 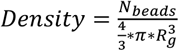 where *R*_*g*_ is radius of gyration of the cluster and *N*_*beads*_ is the total number of beads (stickers + spacers) present in the cluster.

**Figure S5:**
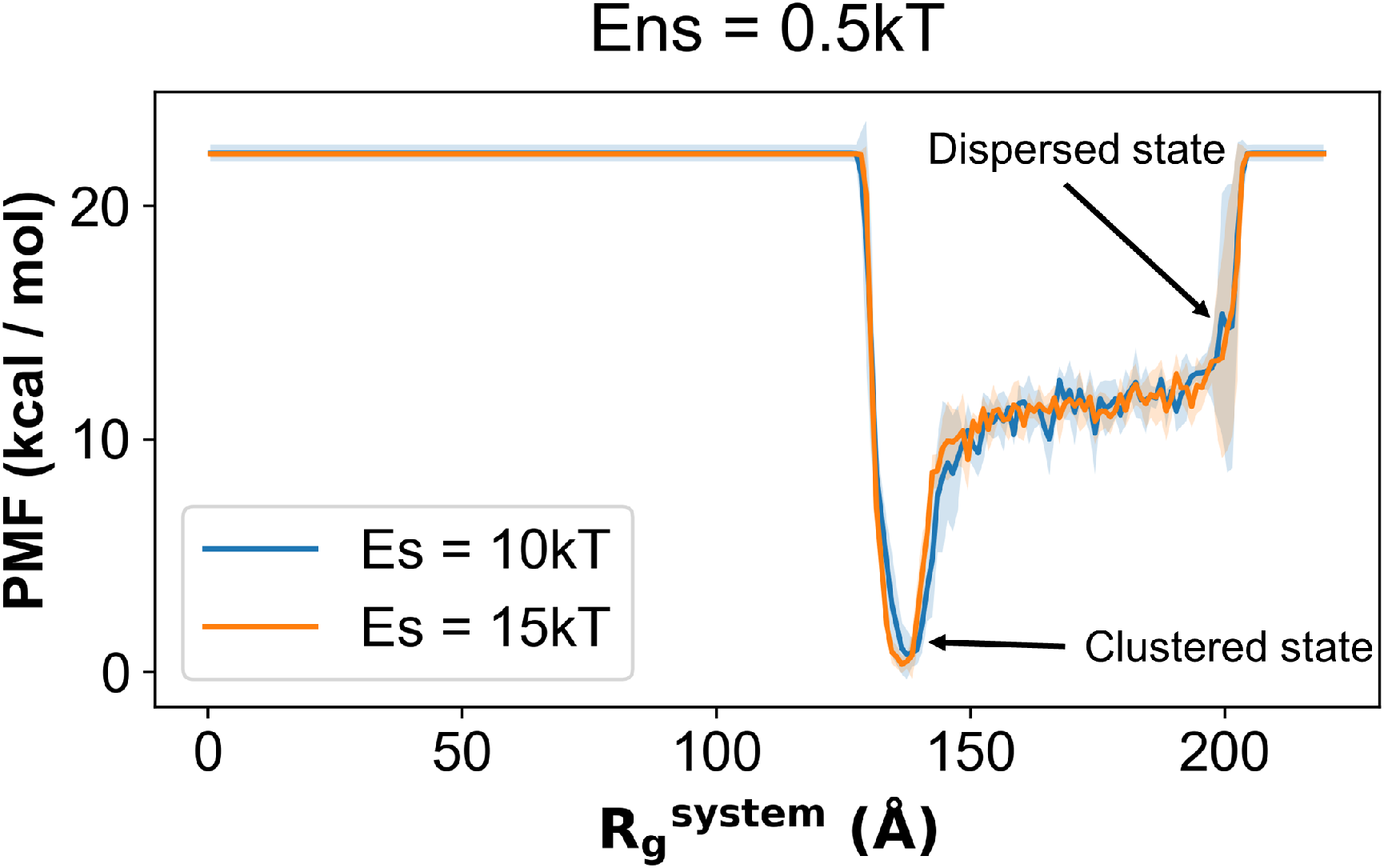
Free energy profile of 400 chains coalescing into one large cluster. The Rg^system^ at the minimum free energy corresponds to the fully clustered state. Each line is an average of 5 stochastic trials. The solid line is the mean and fluctuation envelop represents the standard deviation.

**Figure S6:**
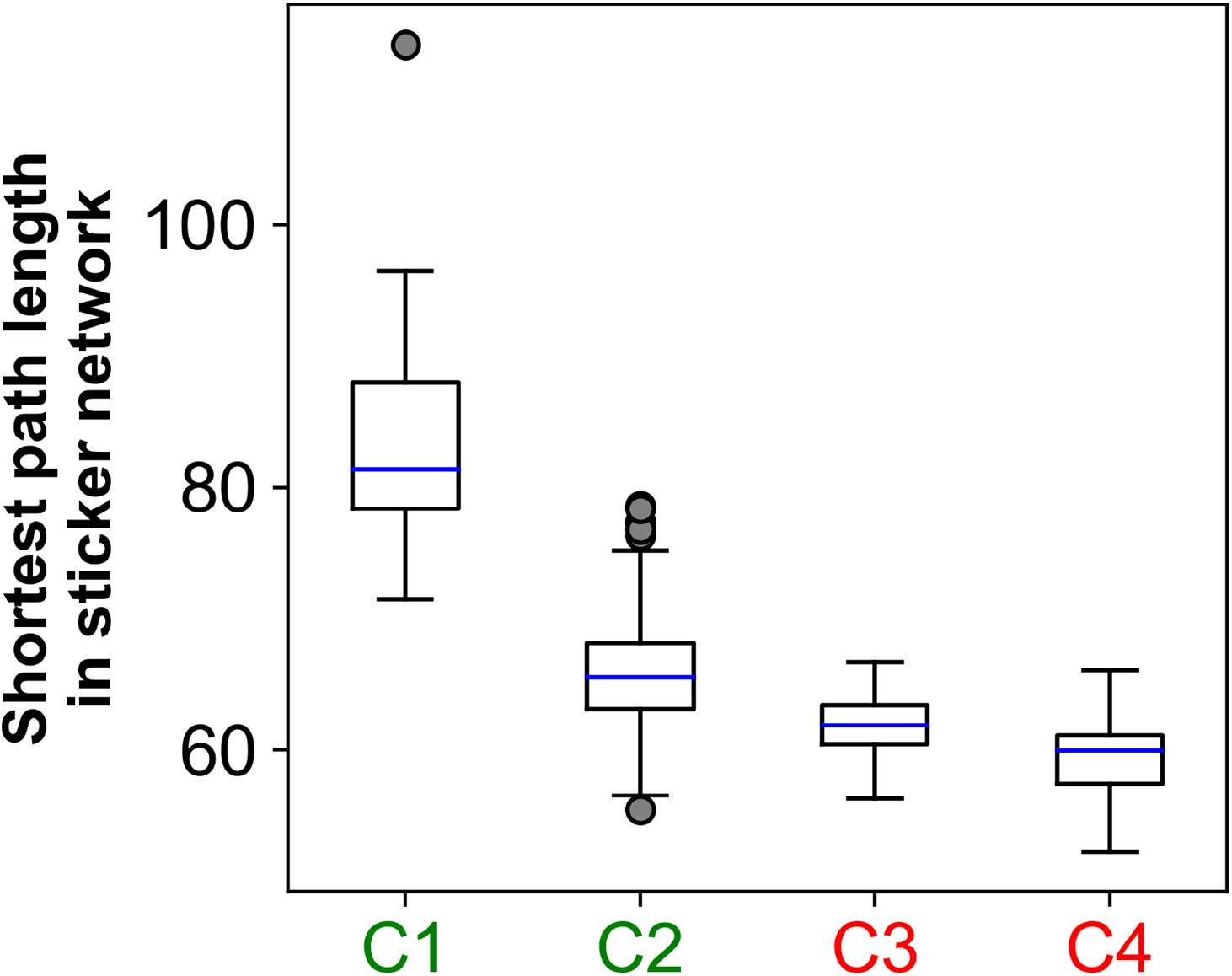
Shortest path length distribution of the sticker-spacer network. Each network (cluster) has 7000 nodes / beads (stickers + spacers). The inter-sticker bonds serve as edges. Starting from one node, the shortest topological paths to all other nodes are computed. Hence, the path length is in the unit of bead count. The color labels are same as in Figure 3. Parameter combinations (Es, Ens) are divided into two categories: C1 (10kT, 0.3kT) and C2 (10kT, 0.5kT) are labelled in green as they undergo fusion; C3 (15kT, 0.3kT) and C4 (15kT, 0.5kT) are labelled in red which do not fuse.

**Figure S7:**
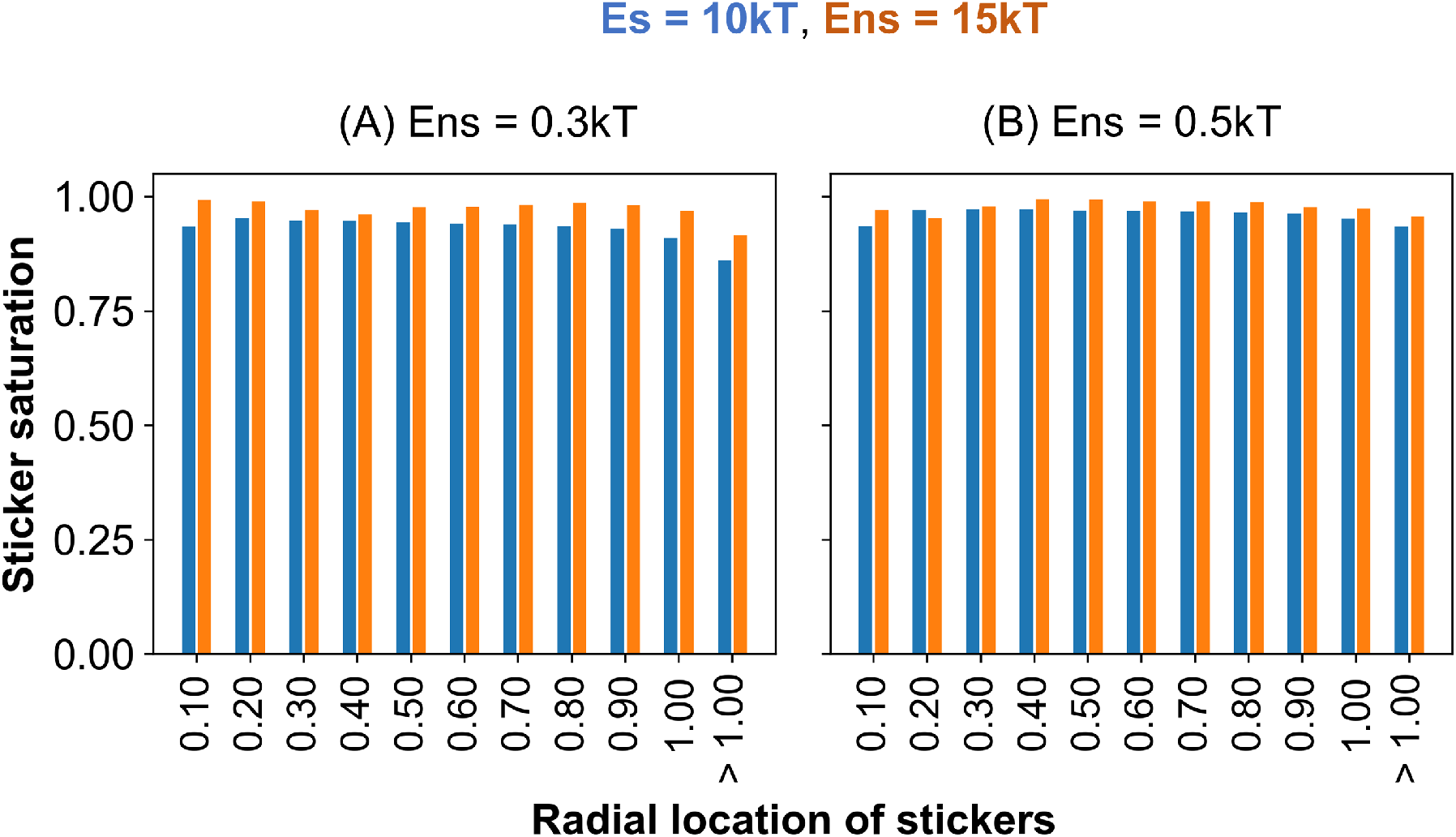
Spatial distribution of saturated stickers within clusters. Extent of sticker saturation as a function of their radial locations within the cluster, at (A) Ens = 0.3kT and (B) Ens = 0.5kT. The blue bars correspond to Es = 10kT, while the orange bars refer to Es = 15kT. For each condition, we compute the distance (*R*_*sticker*_) of each sticker from the cluster center and normalize by cluster radius, *R*_*cluster*_, where 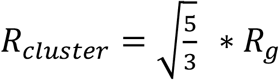 and *R*_*g*_ is the radius of gyration of the cluster. This normalized radial location of stickers is plotted in the horizontal axis. When 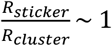, stickers are located near the cluster periphery.

**Figure S8:**
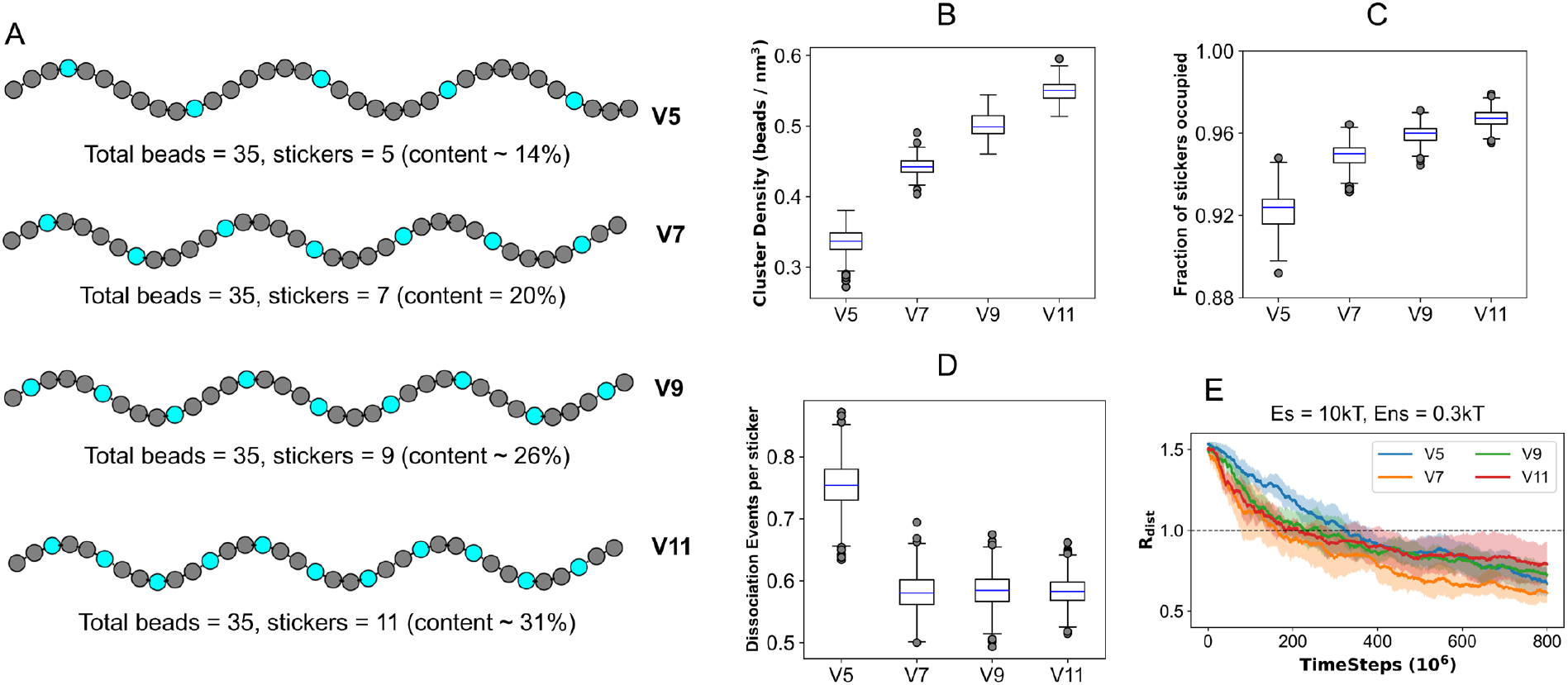
Effect of chain valency on cluster fusion. (A) Illustration of chains with different valencies. Valency of a chain is defined by the number of stickers per chain. Keeping the total beads per chain same, we gradually increase the sticker content. Vn refers to a chain with valency = n. For brevity, only cyan chains are shown; red chains have identical sticker spacer arrangements (similar to Fig. 1A). The interactions remain purely heterotypic, as shown in Fig. 1B. (B) Density of clusters as a function of chain valency. 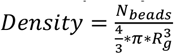 where *R*_*g*_ is radius of gyration of the cluster and *N*_*beads*_ is the total number of beads (stickers + spacers) present in the cluster. (C) Fractional occupancy of stickers that indicates, on average, what fraction of the total sticker population are bonded. (D) Dissociation events refer to how many times an inter-sticker bond is broken in between two successive observation points. Since total sticker counts are different with increasing chain valency, events are normalized by the total stickers present in the respective system. To derive the distributions (B, C, D), for each condition, we run 10 stochastic trials and sample 10 timepoints from each trial; hence each distribution is collected over 100 snapshots. To display the distribution, we used standard boxplot representation or “five-number summary” consisting of the minimum, the maximum, the sample median, and the first and third quartiles. (E) Fusion behavior of clusters as a function of chain valency. Each trajectory is an average over 5 stochastic runs (Solid line: mean, fluctuation envelop: standard deviation). The black dotted line indicates the merging point where two cluster surfaces initiate fusion. *R_dist* is the normalized inter-cluster distance as defined in Fig. 2A. For all these simulations (B-D), Es = 10kT and Ens = 0.3kT.

**Figure S9:**
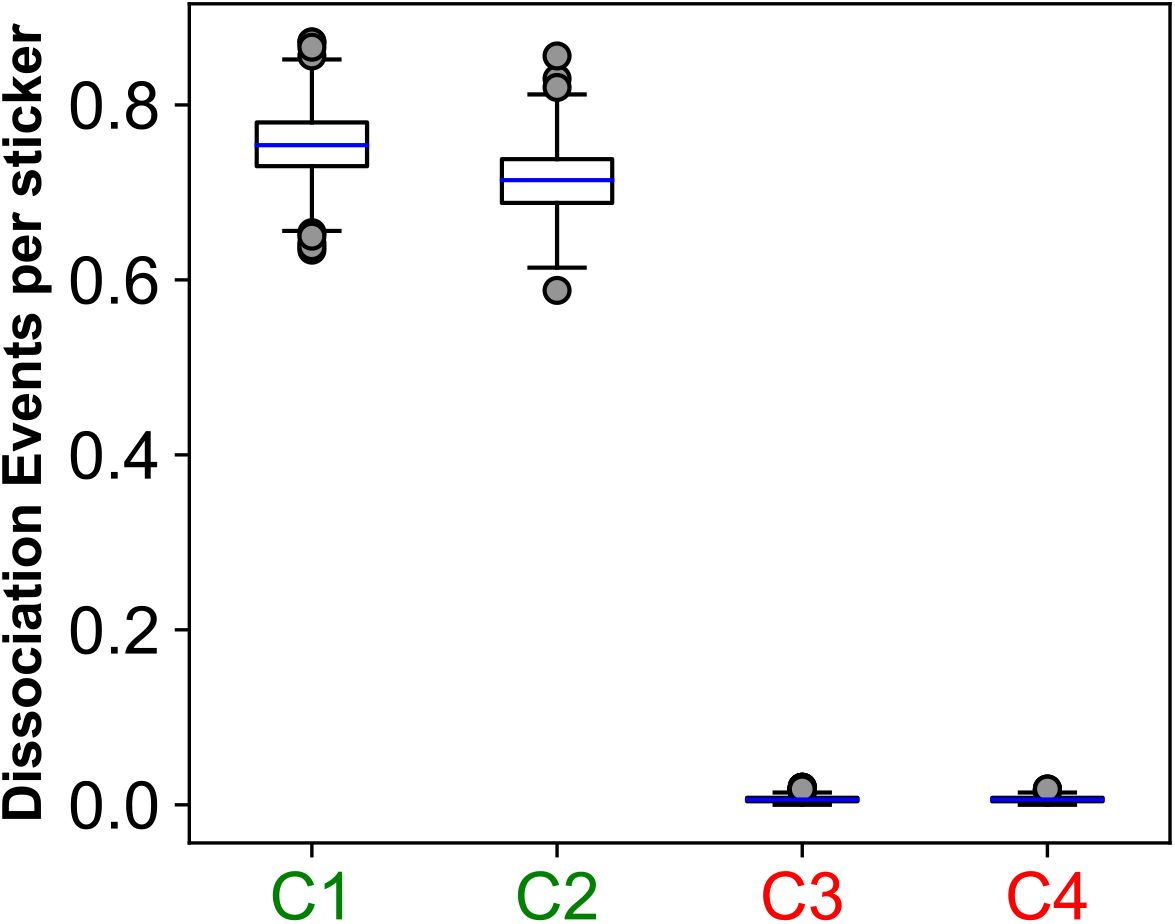
Normalized dissociation kinetics. This is same data as displayed in Fig. 3C. To compare with Fig. S8, inter-sticker dissociation events are normalized by the sticker count in the system. As described earlier, dissociation events refer to how many times an inter-sticker bond is broken in between two successive observation points. Parameter combinations (Es, Ens) are divided into two categories: C1 (10kT, 0.3kT) and C2 (10kT, 0.5kT) are labelled in green as they undergo fusion; C3 (15kT, 0.3kT) and C4 (15kT, 0.5kT) are labelled in red which do not fuse.

**Figure S10:**
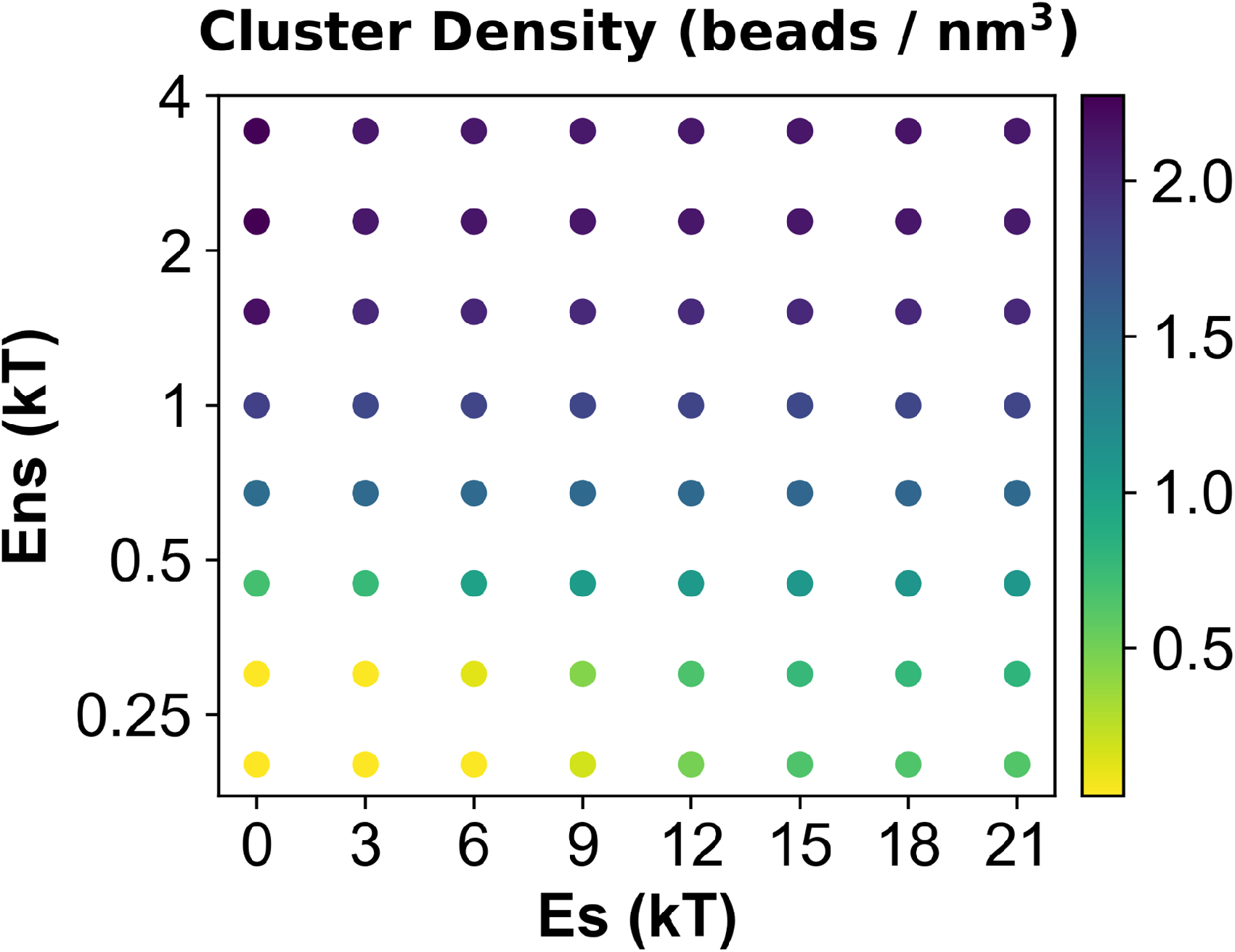
Quantification of cluster density across the phase diagram. Density is computed at the last timepoint of relaxation trajectory. 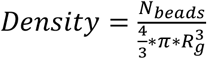 where *R*_*g*_ is radius of gyration of the cluster and *N*_*beads*_ is the total number of beads (stickers + spacers) present in the cluster.

**Figure S11:**
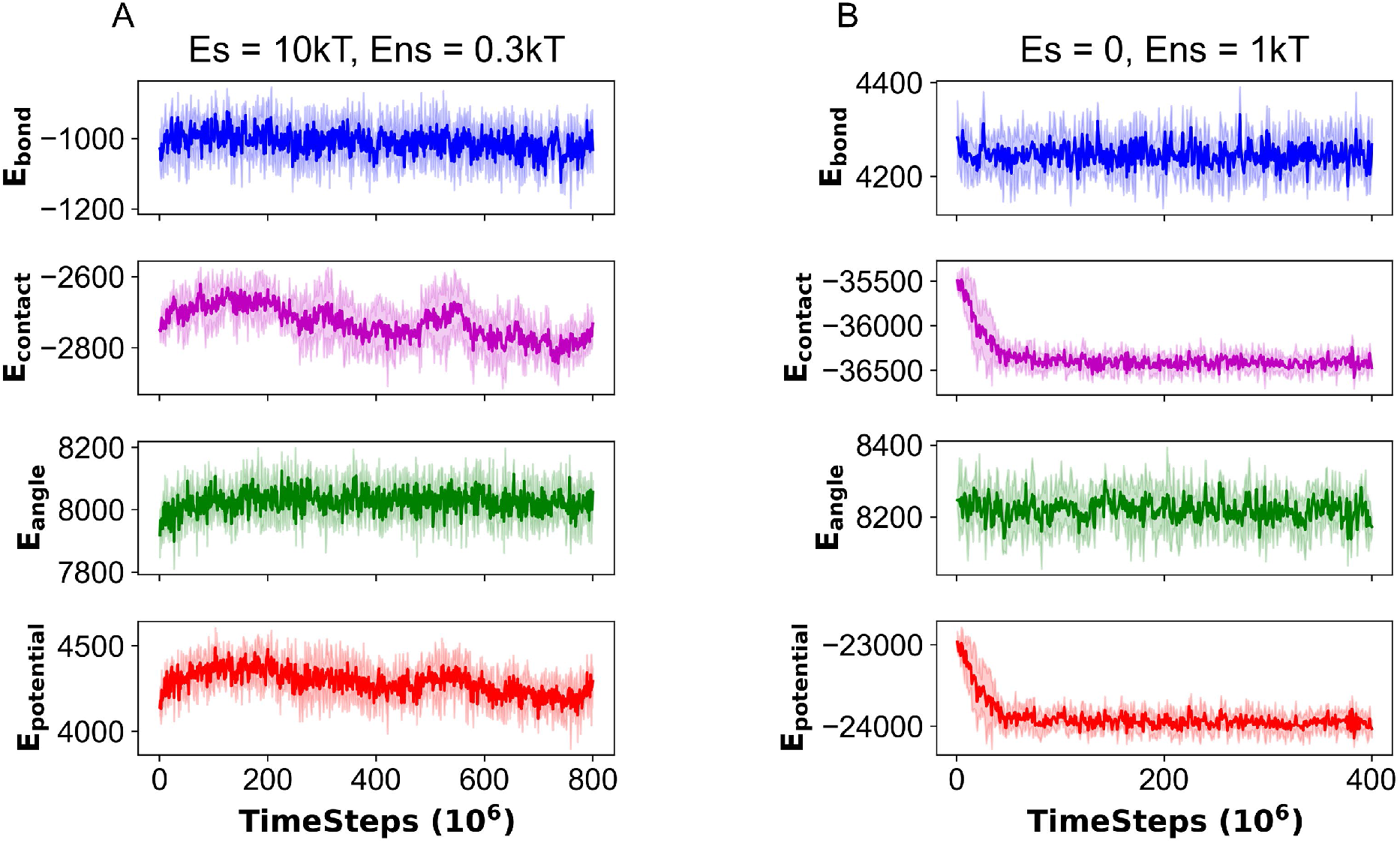
Comparison of energy profiles during fusion of two clusters. made of (A) Heteropolymers (B) Homopolymers. E_bond_ includes all the bonds (permanent and breakable) present in the system. E_contact_ refers to the sum of contact energies coming from the pairwise Lennard-Jones (non-specific) interactions. E_angle_ is angular energy. E_potential_ = E_bond_ + E_pair_ + E_angle_. Energy unit is kcal/mol.

**Figure S12:**
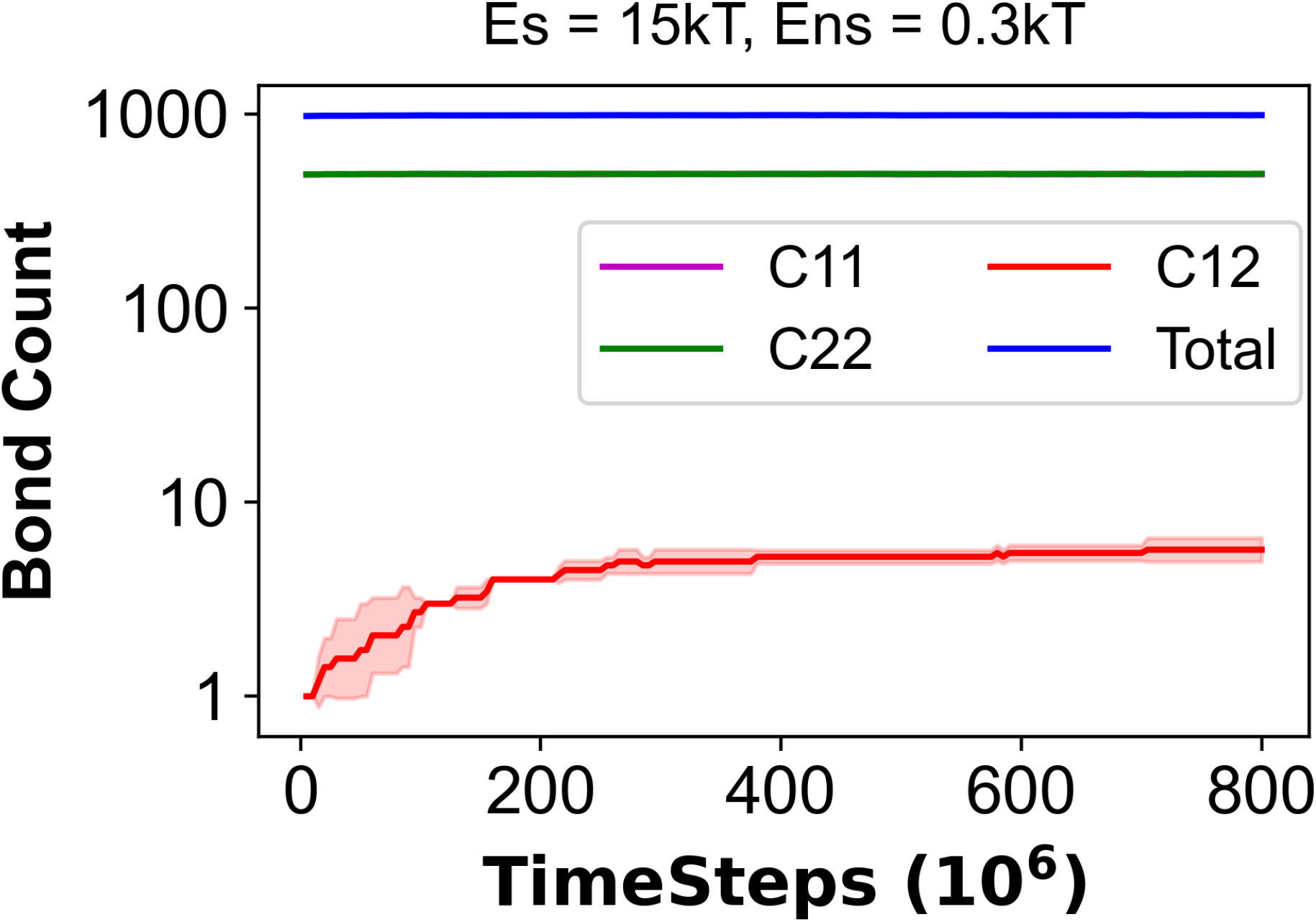
Lack of bond exchange triggers kinetic arrest of sticker-saturated clusters. C11, C22 and C12 indicate intra-cluster-1, intra-cluster-2 and inter-cluster, respectively. “Total” indicates the entire system.

**Figure S13:**
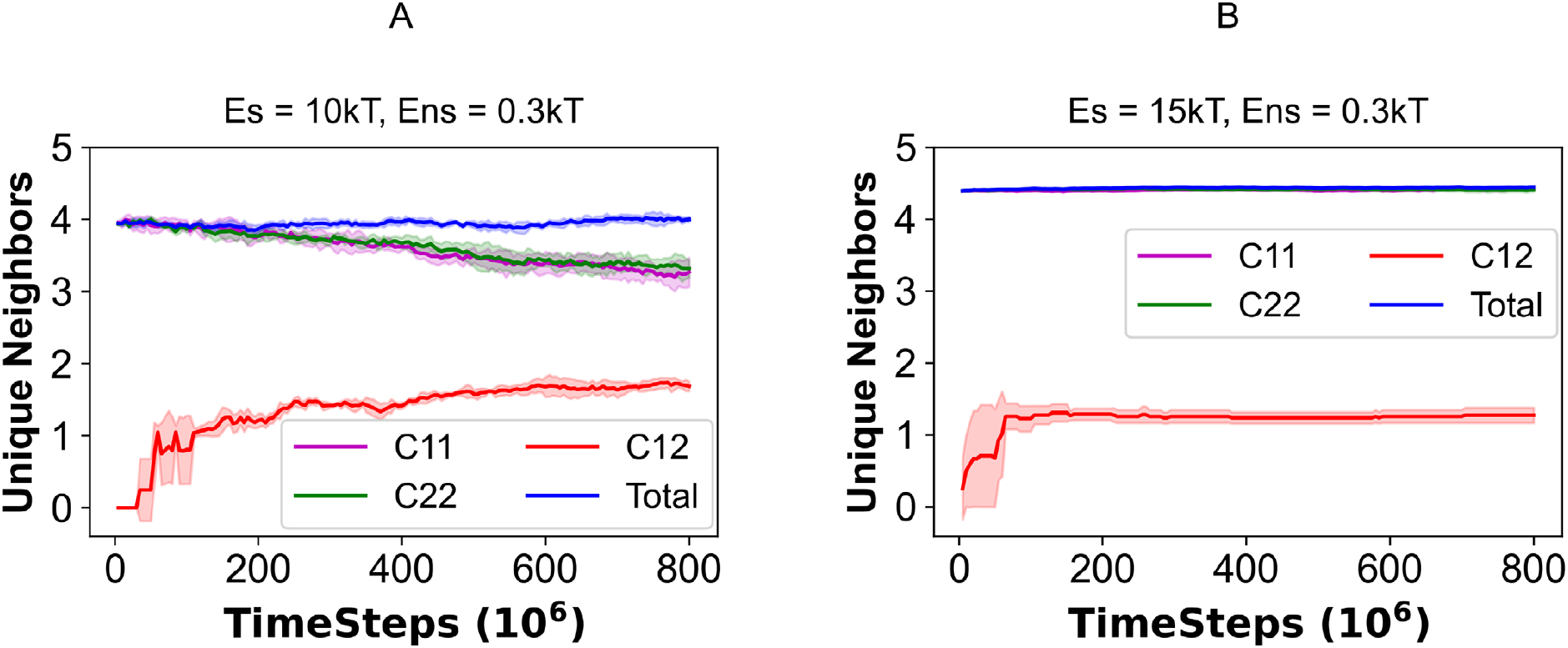
Neighbor exchange during cluster fusion. Since a chain contains 5 stickers, it can be bonded with one neighboring chain at minimum and five neighbors at maximum. If two chains establish two bonds between them, they still have one unique neighbor each. A free chain has no neighbor. Two energy combinations are shown when clusters (A) fuse and (B) do not fuse. C11, C22 and C12 stand for intra-cluster-1, intra-cluster-2 and inter-cluster, respectively. “Total” indicates the entire system.

**Figure S14:**
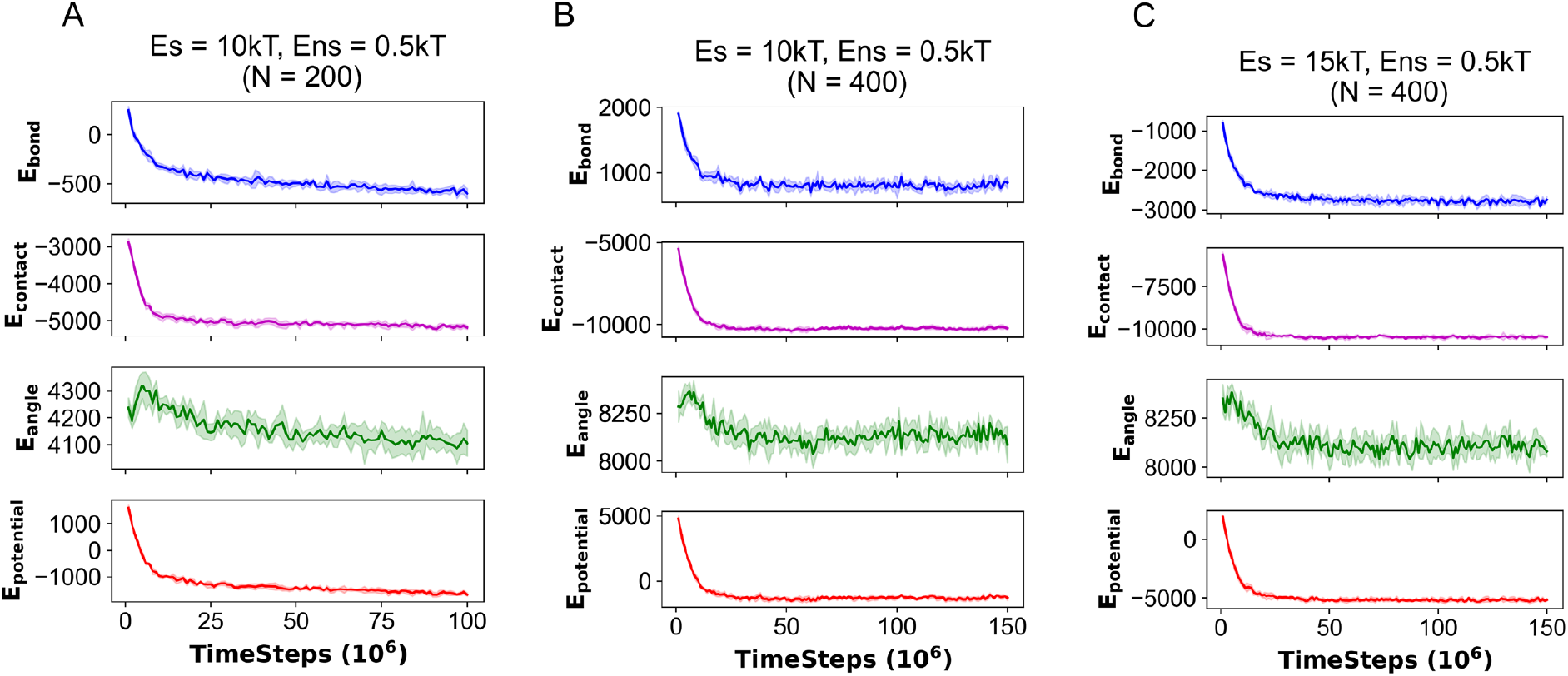
Energy profiles during dispersed to clustered phase transition for sticker-spacer polymers. (A) 200 chains coalescing into one large cluster at Es = 10kT, Ens = 0.5kT. (B, C) 400 chains coalescing into one large cluster at Es = 10kT and 15kT (Ens = 0.5kT) respectively. E_bond_ includes all the bonds (permanent and breakable) present in the system. E_contact_ refers to the sum of contact energies coming from the pairwise Lennard-Jones (non-specific) interactions. E_angle_ is angular energy. E_potential_ = E_bond_ + E_pair_ + E_angle_. Energy unit is kcal/mol. Each trajectory is an average over 5 stochastic runs (Solid line: mean, fluctuation envelop: standard deviation).

**Figure S15:**
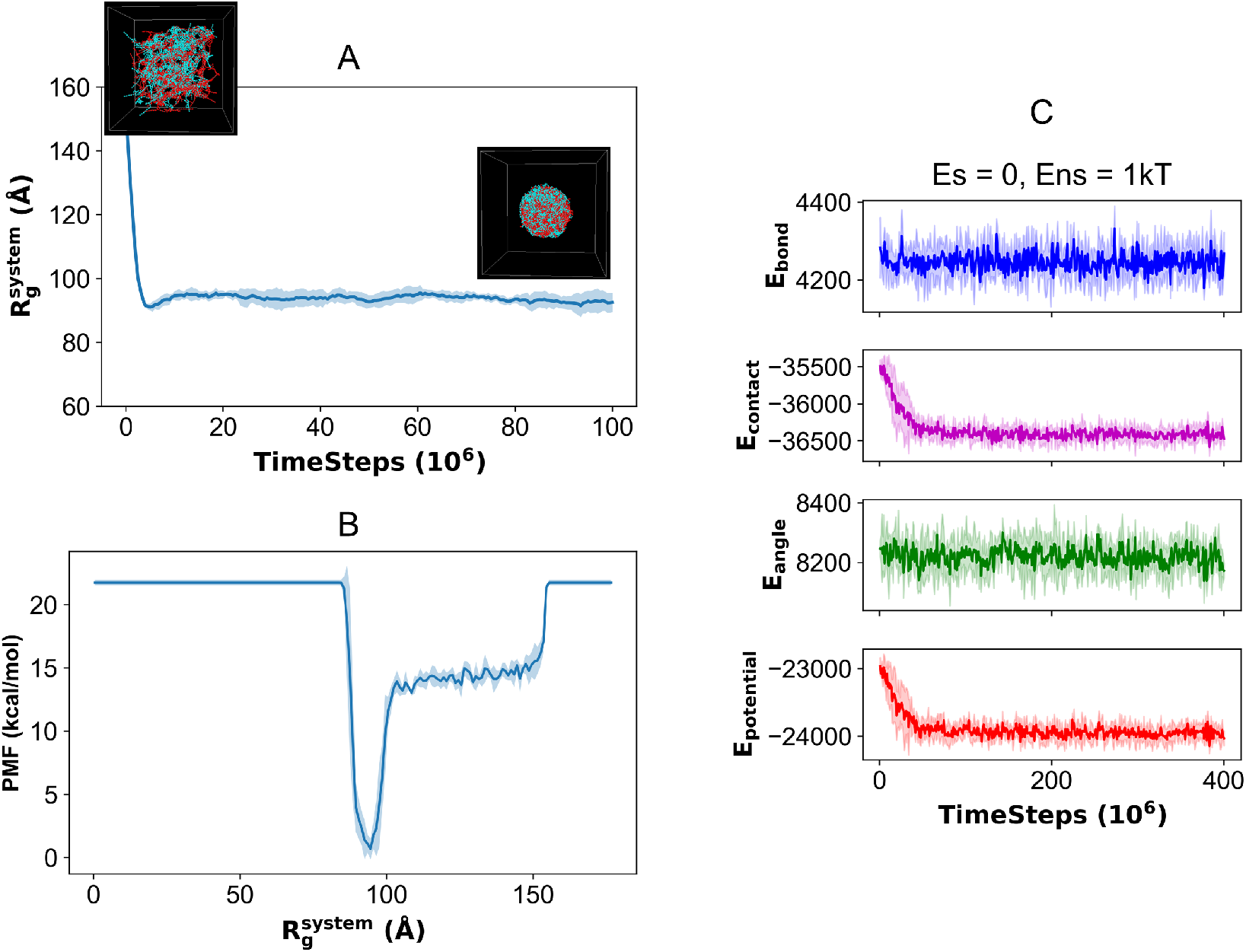
Clustering dynamics of homopolymers (Es = 0, Ens = 1kT). (A) Timecourse of the metadynamics order parameter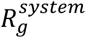, as defined in Figure 1. Insets show first and last timeframes depicting dispersed and clustered states. Since Es = 0, all the beads are colored either red or cyan to indicate that all of them interact in a similar manner. We note that there is no distinction between the chain types here since Es = 0. We still use two color labels to be consistent with Figure 1 color scheme. Free energy profile where the minimum corresponds to the fully clustered state. (C) Energy profile. E_bond_ includes all the bonds (permanent and breakable) present in the system. E_contact_ refers to the sum of contact energies coming from the pairwise Lennard-Jones (non-specific) interactions. E_angle_ is angular energy. E_potential_ = E_bond_ + E_pair_ + E_angle_. Energy unit is kcal/mol. Each trajectory is an average over 5 stochastic runs (Solid line: mean, fluctuation envelop: standard deviation).

## References

1. Banani SF, Lee HO, Hyman AA, Rosen MK. Biomolecular condensates: organizers of cellular biochemistry. Nat Rev Mol Cell Biol. 2017;18(5):285–98.

2. Shin Y, Brangwynne CP. Liquid phase condensation in cell physiology and disease. Science (New York, NY). 2017;357(6357).

3. Lyon AS, Peeples WB, Rosen MK. A framework for understanding the functions of biomolecular condensates across scales. Nat Rev Mol Cell Biol. 2021;22(3):215–35.

4. Mathieu C, Pappu RV, Taylor JP. Beyond aggregation: Pathological phase transitions in neurodegenerative disease. Science (New York, NY). 2020;370(6512):56.

5. Wang B, Zhang L, Dai T, Qin Z, Lu H, Zhang L, et al. Liquid-liquid phase separation in human health and diseases. Signal transduction and targeted therapy. 2021;6(1):290.

6. Choi J-M, Holehouse AS, Pappu RV. Physical Principles Underlying the Complex Biology of Intracellular Phase Transitions. Annual review of biophysics. 2020;49(1):107–33.

7. Mittag T, Pappu RV. A conceptual framework for understanding phase separation and addressing open questions and challenges. Mol Cell. 2022;82(12):2201–14.

8. Ranganathan S, Shakhnovich EI. Dynamic metastable long-living droplets formed by sticker-spacer proteins. Elife. 2020;9.

9. Semenov AN, Rubinstein M. Thermoreversible Gelation in Solutions of Associative Polymers. 1. Statics. Macromolecules. 1998;31(4):1373–85.

10. Choi J-M, Dar F, Pappu RV. LASSI: A lattice model for simulating phase transitions of multivalent proteins. PLOS Computational Biology. 2019;15(10):e1007028.

11. Li P, Banjade S, Cheng H-C, Kim S, Chen B, Guo L, et al. Phase transitions in the assembly of multivalent signalling proteins. Nature. 2012;483(7389):336–40.

12. Harmon TS, Holehouse AS, Rosen MK, Pappu RV. Intrinsically disordered linkers determine the interplay between phase separation and gelation in multivalent proteins. Elife. 2017;6.

13. Martin EW, Holehouse AS, Peran I, Farag M, Incicco JJ, Bremer A, et al. Valence and patterning of aromatic residues determine the phase behavior of prion-like domains. Science (New York, NY). 2020;367(6478):694–9.

14. Li DT, Rudnicki PE, Qin J. Distribution Cutoff for Clusters near the Gel Point. ACS Polymers Au. 2022;2(5):361–70.

15. Flory PJ. Thermodynamics of high polymer solutions. The Journal of chemical physics. 1942;10(1):51–61.

16. Huggins ML. Some properties of solutions of long-chain compounds. The Journal of Physical Chemistry. 1942;46(1):151–8.

17. Zwicker D, Hyman AA, Jülicher F. Suppression of Ostwald ripening in active emulsions. Physical review E, Statistical, nonlinear, and soft matter physics. 2015;92(1):012317.

18. Söding J, Zwicker D, Sohrabi-Jahromi S, Boehning M, Kirschbaum J. Mechanisms for Active Regulation of Biomolecular Condensates. Trends in cell biology. 2020;30(1):4–14.

19. Nakashima KK, van Haren MHI, André AAM, Robu I, Spruijt E. Active coacervate droplets are protocells that grow and resist Ostwald ripening. Nat Commun. 2021;12(1):3819.

20. Cuylen S, Blaukopf C, Politi AZ, Müller-Reichert T, Neumann B, Poser I, et al. Ki-67 acts as a biological surfactant to disperse mitotic chromosomes. Nature. 2016;535(7611):308–12.

21. Wang Z, Yang C, Guan D, Li J, Zhang H. Cellular proteins act as surfactants to control the interfacial behavior and function of biological condensates. Developmental cell. 2023;58(11):919-32.e5.

22. Folkmann AW, Putnam A, Lee CF, Seydoux G. Regulation of biomolecular condensates by interfacial protein clusters. Science (New York, NY). 2021;373(6560):1218–24.

23. Erkamp NA, Sneideris T, Ausserwöger H, Qian D, Qamar S, Nixon-Abell J, et al. Spatially nonuniform condensates emerge from dynamically arrested phase separation. Nature Communications. 2023;14(1):684.

24. Snead WT, Jalihal AP, Gerbich TM, Seim I, Hu Z, Gladfelter AS. Membrane surfaces regulate assembly of ribonucleoprotein condensates. Nat Cell Biol. 2022;24(4):461–70.

25. Lee DSW, Choi C-H, Sanders DW, Beckers L, Riback JA, Brangwynne CP, et al. Size distributions of intracellular condensates reflect competition between coalescence and nucleation. Nature Physics. 2023;19(4):586–96.

26. Jan Bachmann S, Petitzon M, Mognetti BM. Bond formation kinetics affects self-assembly directed by ligand–receptor interactions. Soft Matter. 2016;12(47):9585–92.

27. Xiang YX, Shan Y, Lei QL, Ren CL, Ma YQ. Dynamics of protein condensates in weak-binding regime. Phys Rev E. 2022;106(4-1):044403.

28. Ronceray P, Zhang Y, Liu X, Wingreen NS. Stoichiometry Controls the Dynamics of Liquid Condensates of Associative Proteins. Phys Rev Lett. 2022;128(3):038102.

29. Garaizar A, Espinosa JR, Joseph JA, Collepardo-Guevara R. Kinetic interplay between droplet maturation and coalescence modulates shape of aged protein condensates. Scientific Reports. 2022;12(1):4390.

30. Ranganathan S, Shakhnovich E. The physics of liquid-to-solid transitions in multi-domain protein condensates. Biophys J. 2022;121(14):2751–66.

31. Banani SF, Rice AM, Peeples WB, Lin Y, Jain S, Parker R, et al. Compositional Control of Phase-Separated Cellular Bodies. Cell. 2016;166(3):651–63.

32. Lin AZ, Ruff KM, Dar F, Jalihal A, King MR, Lalmansingh JM, et al. Dynamical control enables the formation of demixed biomolecular condensates. Nature Communications. 2023;14(1):7678.

33. Thompson AP, Aktulga HM, Berger R, Bolintineanu DS, Brown WM, Crozier PS, et al. LAMMPS - a flexible simulation tool for particle-based materials modeling at the atomic, meso, and continuum scales. Computer Physics Communications. 2022;271:108171.

34. Plimpton S. Fast Parallel Algorithms for Short-Range Molecular Dynamics. Journal of Computational Physics. 1995;117(1):1–19.

35. Laio A, Gervasio FL. Metadynamics: a method to simulate rare events and reconstruct the free energy in biophysics, chemistry and material science. Reports on Progress in Physics. 2008;71(12):126601.

36. Barducci A, Bussi G, Parrinello M. Well-Tempered Metadynamics: A Smoothly Converging and Tunable Free-Energy Method. Physical Review Letters. 2008;100(2):020603.

37. Ranganathan S, Dasmeh P, Furniss S, Shakhnovich E. Phosphorylation sites are evolutionary checkpoints against liquid–solid transition in protein condensates. Proceedings of the National Academy of Sciences. 2023;120(20):e2215828120.

38. Jewett AI, Stelter D, Lambert J, Saladi SM, Roscioni OM, Ricci M, et al. Moltemplate: A Tool for Coarse-Grained Modeling of Complex Biological Matter and Soft Condensed Matter Physics. Journal of molecular biology. 2021;433(11):166841.

39. Martínez L, Andrade R, Birgin EG, Martínez JM. PACKMOL: a package for building initial configurations for molecular dynamics simulations. J Comput Chem. 2009;30(13):2157–64.

40. Fiorin G, Klein ML, Henin J. Using collective variables to drive molecular dynamics simulations. Mol Phys. 2013;111(22-23):3345–62.

41. de Buyl P, Nies E. A parallel algorithm for step- and chain-growth polymerization in molecular dynamics. J Chem Phys. 2015;142(13):134102.

42. Stukowski A. Visualization and analysis of atomistic simulation data with OVITO–the Open Visualization Tool. Modelling and Simulation in Materials Science and Engineering. 2010;18(1):015012.

43. Jawerth L, Fischer-Friedrich E, Saha S, Wang J, Franzmann T, Zhang X, et al. Protein condensates as aging Maxwell fluids. Science (New York, NY). 2020;370(6522):1317–23.

44. Wang H, Kelley FM, Milovanovic D, Schuster BS, Shi Z. Surface tension and viscosity of protein condensates quantified by micropipette aspiration. Biophys Rep (N Y). 2021;1(1).

45. Lazar T, Tantos A, Tompa P, Schad E. Intrinsic protein disorder uncouples affinity from binding specificity. Protein Sci. 2022;31(11):e4455.

46. Michieletto D, Marenda M. Rheology and Viscoelasticity of Proteins and Nucleic Acids Condensates. JACS Au. 2022;2(7):1506–21.

47. Alshareedah I, Kaur T, Banerjee PR. Methods for characterizing the material properties of biomolecular condensates. Methods in enzymology. 646: Elsevier; 2021. p. 143–83.

48. Alshareedah I, Borcherds WM, Cohen SR, Singh A, Posey AE, Farag M, et al. Sequence-specific interactions determine viscoelasticity and ageing dynamics of protein condensates. Nature Physics. 2024.

